# Overcoming Extrapolation Challenges of Deep Learning by Incorporating Physics in Protein Sequence-Function Modeling

**DOI:** 10.1101/2025.11.09.687530

**Authors:** Shrishti Barethiya, Jian Huang, Xiao Liu, Hui Guan, Jianhan Chen

## Abstract

Understanding protein sequence-to-function relationship is crucial to assist studies of genetic diseases, protein evolution, and protein engineering. The sequence-to-function relationship of proteins is inherently complex due to multi-site high-dimensional correlation and structural dynamics. Deep learning algorithms such as (graph) convolutional neural networks and recently transformers have become very popular for learning the protein sequence-to-function mapping from deep mutational scanning data and available structures. However, it remains very challenging for these models to achieve accurate extrapolation when predicting functional effect of variants with positions or mutation types not seen in the training data. We propose that incorporating the physics of protein interactions and dynamics can be an effective approach to overcome the extrapolation limitations. Specifically, we demonstrate that physics-based modeling can be used to quantify the energetic effects of mutations and that incorporating these physical energetics directly within the convolution and graph convolution neural networks can significantly improve the performance of positional and mutational extrapolation compared to models without biophysics-inspired features. Our results support the effectiveness of leveraging physical knowledge in overcoming the limitation of data scarcity.

## Introduction

Proteins play crucial and diverse roles in nearly all biological processes. The function of a specific protein is ultimately dictated by its amino acid sequence. The physicochemical properties of amino acids, their spatial arrangements as defined by three-dimensional structures, and conformational dynamics together provide the physical basis of protein biological functions. Understanding the protein sequence-to-function relationship, that is, how sequence governs their biological functions, is a fundamental problem central to the studies of protein evolution, diagnosis of genetic disease, and protein engineering and design [1–4]. While homology-based annotation of protein function given the sequence has been highly successful [5–7], the sequence-to-function mapping remains inherently challenging. This is especially true in the prediction of functional effects of one or a few specific mutations of a given protein, due to the complex high-dimensional correlation between sequence and function that arise from frequently nontrivial impacts on physiochemical, structural and dynamic properties of the protein.

In the past couple of decades, advancements in high-throughput functional assays, especially deep mutational scanning or DMS [8–12], have generated large sets of functional readouts in the sequence space of protein variants. DMS integrates large genetic mutant libraries, high-throughput phenotyping, and DNA sequencing of both input and post-selection cells, enabling examination of functional impact of up to tens of thousands of mutants in a systematic and economical fashion [13–15]. Each step in a DMS experiment may have inherent limitations. For example, the library construction technique can introduce mutation bias, off-target issues, or efficiency constraints, and, most critically, it is typically limited to handling up to tens of thousands of variants [16,17]. Since the sequence space grows exponentially with length, achieving comprehensive DMS analyses of large proteins becomes prohibitively expensive and is generally infeasible. Leveraging data from DMS experiments, genome-wide sequencing [18], and vast computational and experimental structural databases [19–22], machine learning (ML) models have been trained to make quantitative predictions of sequence-to-function molecular phenotypes [23–29]. These predictors, also commonly referred to as variant effect predictors (VEPs), provide powerful tools for many applications, including designing a protein variant optimizing functions of naturally occurring proteins [30], creating protein variants of exceptional fitness [31,32], and developing treatment strategies of genetic diseases [33–36].

VEPs can be broadly categorized into unsupervised and supervised learning models. These models generally utilize a combination of four major input feature types: protein wildtype or mutant sequence, amino acid properties, protein structure, and evolutionary information [27]. Unsupervised learning models typically exploit extensive protein sequence data and multi sequence alignment (MSA) to learn complex epistatic interactions or constraints between sites. For example, DeepSequence [37] is a Bayesian deep latent-variable model trained on large DMS datasets of proteins and RNA domains. This model learns the probability distribution of sequences, predicts mutational effects, and models the evolutionary fitness landscape within families. SeqDesign [38] is an alignment-free autoregressive generative model that uses dilated convolutional neural networks trained on protein family sequences, achieving similar prediction accuracies benchmarked on 40 protein DMS datasets. SeqDesign offers major improvements where MSA is not robust, such as predicting effects of indels, disordered proteins, and designing proteins like antibodies.

Supervised learning VEPs such as Envision [25], positive-unlabeled learning [39], and ECNet [40] are trained on experimental mutagenesis/DMS datasets, using approaches ranging from decision trees to protein language models. In particular, Gelman et al [41] have designed the NN4dms deep learning framework that includes a set of amino acid properties (AAIndex [42]) features and/or 3D structures in addition to sequences, and evaluated four ML algorithms ranging from linear regression (LR), fully connected neural network (NN), sequence convolutional neural network (CNN), to graph convolutional neural network (GCN). All four models performed well with DMS datasets containing single and higher-order mutations compared to biophysics-based and unsupervised methods, especially with CNN and GCN. A key lesson is that the performance depends critically on the sequence coverage. This is particularly evident when challenged with mutational and positional extrapolations [41,43]. In mutational extrapolation, models aim to predict the effects of mutation types not observed at the site of interest during training; in positional extrapolation, models aim to predict the effects of mutations at entirely unseen sites. All NN4dms models struggle in mutational extrapolation and largely fail in positional extrapolation in all testing cases [41]. These results highlight arguably the most common bottleneck in training supervised VEPs, the data scarcity problem, especially for larger proteins where comprehensive coverage in DMS experiments becomes infeasible.

Novel information must be included to supplement DMS datasets to overcome the data scarcity problem in general. One elegant strategy is to use pre-trained models, such as large language models (LLMs) derived from sequence databases. For example, the geometric vector perceptron multi-sequence alignment (GVP-MSA) model [32] utilizes pre-trained MSA transformer model [44] to encode higher-order sequence dependencies, which is then integrated into the GVP-graph neural network alongside extracted structural features. The model has shown to perform well for positional extrapolation of single-variant effects and predicting the higher-order mutational effect from datasets containing only single mutations. Another strategy is to incorporate fundamental biophysics of protein structure, dynamics and interactions during model training [45–47]. The premise here is that these physical principles ultimately determine how any sequence variation may affect function and are thus ideally suited for overcoming extrapolation limitations. In a recent mutational effect transfer learning (METL) model [48], a transformer-based neural network was pre-trained on large-scale synthetic data derived from Rosetta-based molecular modeling [49] and then fine-tuned on the experimental DMS data. The authors evaluated two types of models, METL-local and METL-global, the first of which is protein-specific and the latter is trained on a broad range of proteins [48]. Even though METL-global suffers from severe over-fitting, both models demonstrated superior performance compared to NN4dms when sequence coverage is limited. Especially for positional extrapolation, modest Spearman correlation of ∼0.5 was achieved for all test proteins with either METL-local or METL-global [48]. These successes clearly support the effectiveness of incorporating biophysics to overcome data scarcity in deep learning.

In this work, we developed an efficient and direct strategy for incorporating biophysics-based features to train accurate supervised protein-specific VEPs and to overcome both mutational and positional extrapolation challenges. The strategy builds upon our previous work in predicting the mutational effects of the gating voltage of big potassium (BK) channels [50], where random forest models, trained with less than 500 data points, achieve accurate performance when supplemented with biophysical features including Rosetta-based energy terms and structural dynamics from all-atom simulations. The predictor correctly recapitulates the hydrophobic gating mechanism of BK channels and captures nontrivial voltage gating properties in regions where few mutations are known [50]. Several novel mutations were subsequently confirmed by experiments. The workflow presented in this work only requires evaluation of Rosetta energies for all 19 possible single mutations for each residue site of a given protein, each of which involves a short optimization of side chain repacking within the vicinity of the mutation. The resulting Rosetta energy terms are used directly to quantify the impacts of any single or multiple variants on various types of physical interactions, assuming additivity. The resulting CNN and GCN models are comparable to those trained without biophysical features in NN4dms for random splitting but provide superior extrapolation performance comparable to METL. The overall workflow is highly efficient, mainly requiring only sequence length x 19 Rosetta rotamer repacking evaluations, and can be readily extended to any protein or protein complex, allowing an accurate VEP trained within few minutes to hours given the DMS dataset.

## Methods

### Model proteins and DMS datasets

Following NN4dms [41], five protein DMS datasets are considered in this work, including avGFP (*Aequorea victoria* green fluorescent protein) [51], Bgl3 (*β*-glucosidase enzyme) [11], GB1 (IgG-binding domain of protein G) [52], Pab1 (RRM2 domain of the *Saccharomyces cerevisiae* poly(A)- binding protein) [13], and Ube4b (U-box of the murine E3 ligase ubiquitination factor E4B) [53]. These proteins cover sizes from 56 to 501 residues and represent distinct topologies from helical, mixed helix/beta to full beta-barrel (***Figure 1***). Their sequence lengths, functions, and DMS dataset sizes are summarized in ***Table 1***. The functional scores are available publicly for the avGFP, Pab1, and Ube4b, whereas for GB1 and Bgl3, functional scores are processed through Enrich2 [54] by Gitter and Coworkers [41] based on raw sequencing read counts. The dataset consists of substitutions and deletion mutations. Before the model training, all the deletions have been removed from the datasets and the final datasets only contain missense substitutions. The distributions of final DMS data points (***Figure 1***) display signficant heterogeneity along the protein sequences except for GB1 with exceptional number of over half million variants. Most variants in the datasets are single and double mutation variants (***Figure S1A***), with avGFP containing the broadest distribution in number of mutations per variants (up to 15). The higher order variants could inform correlation between different mutation sites and thus potentially benefit VEP models to make more accurate positional and mutational extrapolations. Of note, only 43% and 38% of possible site mutations are represented for larger proteins avGFP and Bgl3, respectively, and many of mutations are only sample as part of high order multi-site mutation variants (***Figure S1B***). Heterogenity of structures, functions, DMS data distributions of these five protein cases provides a good benchmark for model performance assessments.

**Figure 1.**
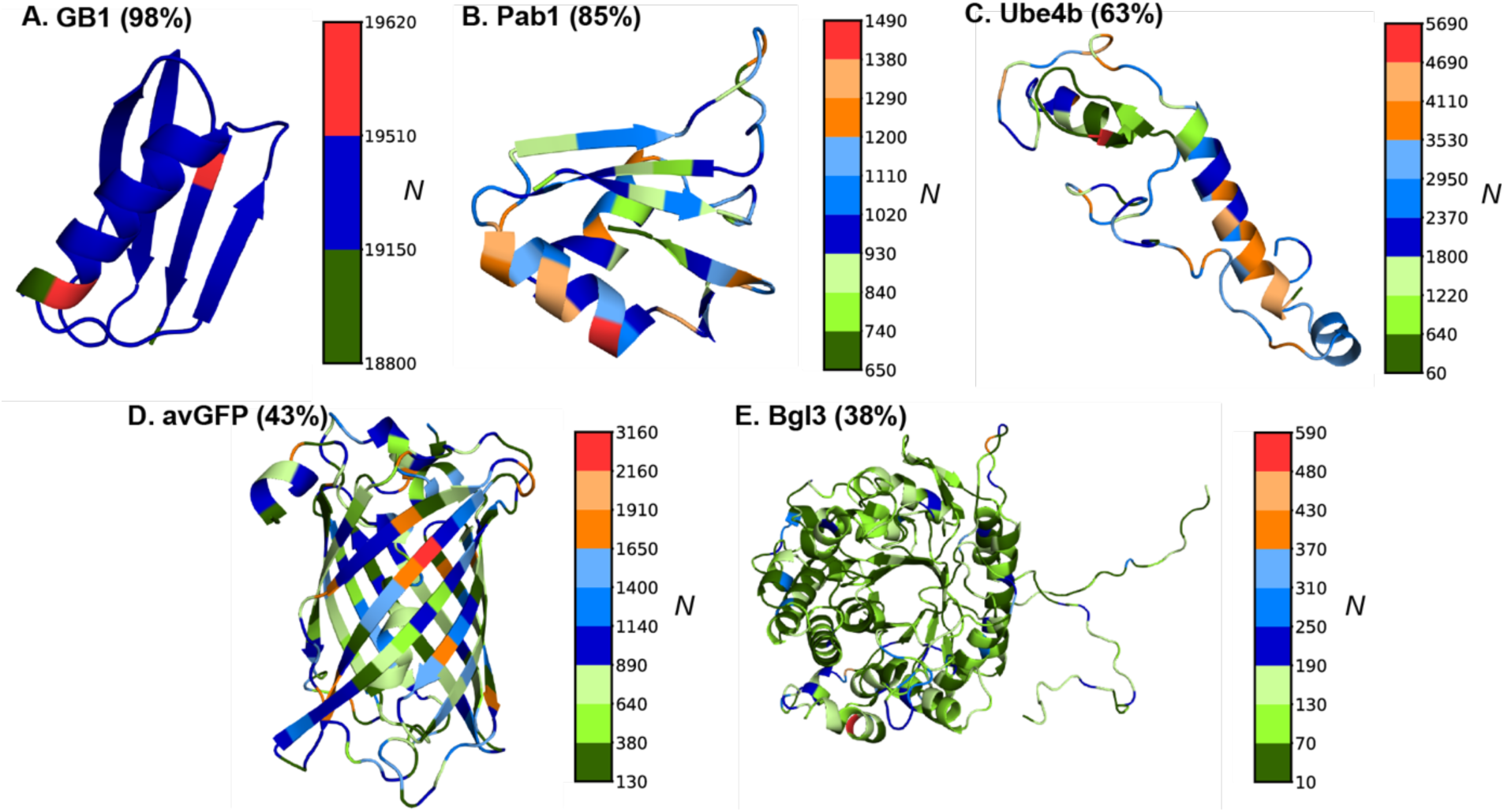
Model protein structures and mutational data point distributions. All proteins are represented using cartoons with PDB structures. The PDB IDs used are: 2QMT (GB1), 1CVJ (Pab1), 2KR4 (Ube4b), 1EMM (avGFP), and 1GNX (Bgl3). Each site on the structure is colored based on the number of DMS variants containing mutations at that site (N). The coverage of unique mutations sampled in the dataset is given in parenthesis next to the protein name.

**Table 1:**
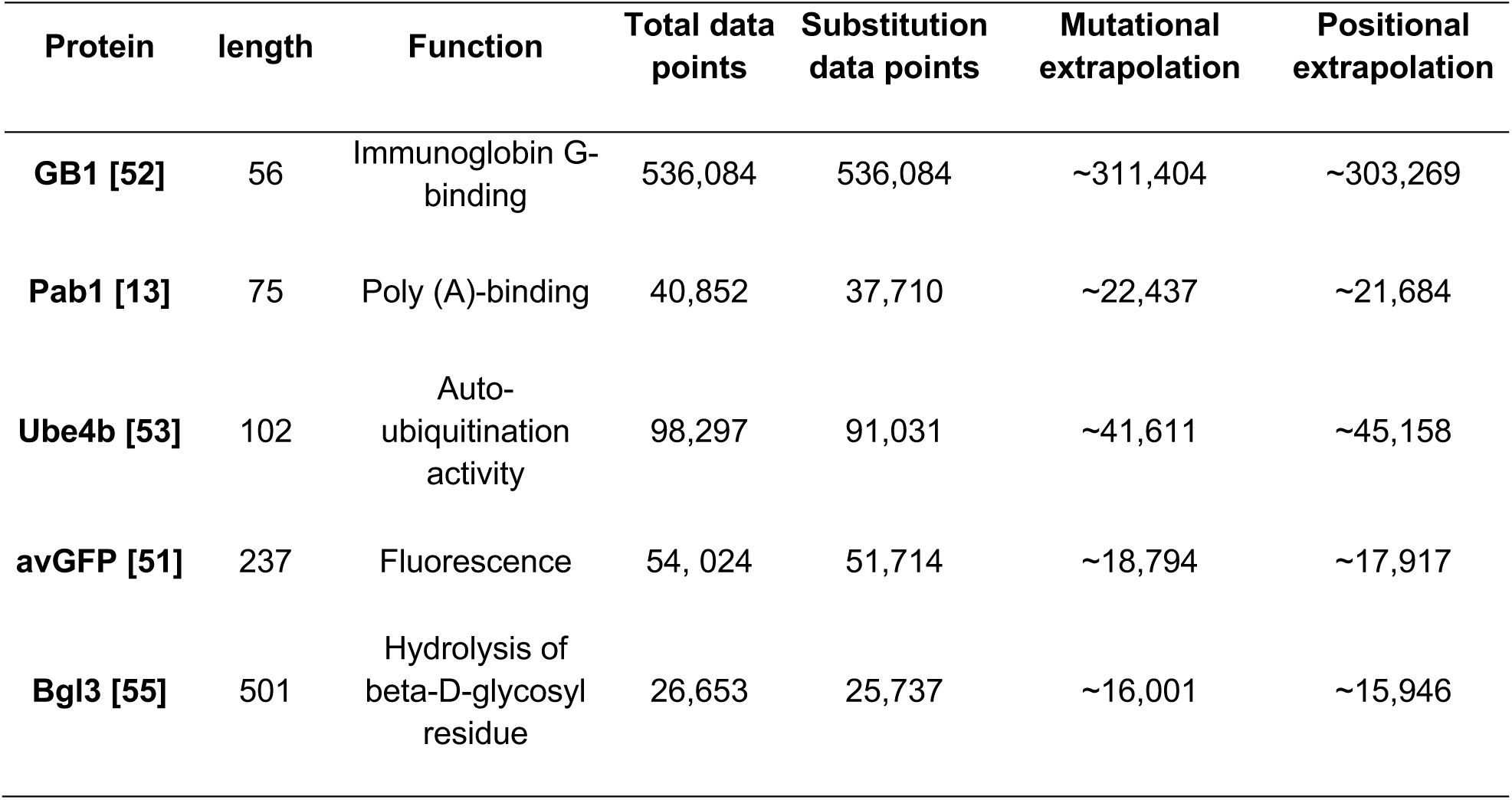
Summary of the five proteins and DMS datasets, ordered in decreasing percentage of sequence space coverage in the dataset.

| Protein | length | Function | Total data points | Substitution data points | Mutational extrapolation | Positional extrapolation |
| --- | --- | --- | --- | --- | --- | --- |
| **GB1 [52]** | 56 | Immunoglobulin G-binding | 536,084 | 536,084 | ~311,404 | ~303,269 |
| **Pab1 [13]** | 75 | Poly (A)-binding | 40,852 | 37,710 | ~22,437 | ~21,684 |
| **Ube4b [53]** | 102 | Auto-ubiquitination activity | 98,297 | 91,031 | ~41,611 | ~45,158 |
| **avGFP [51]** | 237 | Fluorescence | 54, 024 | 51,714 | ~18,794 | ~17,917 |
| **Bgl3 [55]** | 501 | Hydrolysis of beta-D-glycosyl residue | 26,653 | 25,737 | ~16,001 | ~15,946 |

### Calculation of Rosetta energetic features

The Rosetta FastRelax protocol [56] was used to relax the initial PDB structure of each protein for 5 independent runs of 100 iterations. During relaxation, both backbone and side chain degrees of freedom were allowed, and the lowest energy pose from 5 runs was eventually selected as the starting point for the subsequent mutational ΔΔG calculations.

The predict_ddG.py script from the PyRosetta [57] package was used to calculate the free energies of all possible single mutations of each protein. Within the script, side chain substitution is first made at a given site, and the PackRotamerMover method is then used to repack side chains of surrounding residues within a certain cut-off radius using the Dunbrack rotamer libraries and minimize the conformational energy based on the REF2015 energy function [49]. For an input structure, the script will internally calculate the ΔG values (the folding Gibbs free energy versus the fully unfold status) for the wildtype and a mutant with a specified mutation site after repacking and report the final ΔΔG ( = 1′Gwildtype - 1′Gmutant ). To achieve stable energy evaluations and balance the tradeoff between re-packing iterations and computational cost (few seconds per mutation), we focused on optimizing two key parameters for the protocol (see detailed discussion in the Results section): the cutoff radius (repacking radius) and *nloops*. The repacking radius is used to select residues to be repacked by including surrounding residues that have a C_α_ (for glycine) or C_β_ (other residues) atoms within the cutoff radius of the mutated site. The *nloop* parameter defines the side chain repacking iterations before outputting the final score.

We have shown that it is advantageous include individual Rosetta energy components in addition to the total ΔΔG [49]. Even though the Rosetta energy score function consists of 19 energy terms, only 8 terms with substantial variances were selected to be included in training: Lennard-Jones attractive and repulsive between atoms in the same and different residue, Lazaridis-Karplus solvation energy, Coulombic electrostatic potential with distance-dependent dielectric, backbone hydrogen bonds, sidechain-backbone and sidechain-sidechain hydrogen bond energy, disulfide geometry potential, energy term based on Ramachandran maps, probability of amino acid at different backbone torsion and difference in Rosetta free energy of the wildtype (***Table S1***).

### Atomistic molecular dynamics simulations

Short atomistic molecular dynamics (MD) simulations with explicit solvents and ions were conducted for each protein to quantify the inherent flexibility of each residue in the context of the protein structure and its native-like environment, as characterized by the root mean square fluctuation (RMSF) of the Cα atom. The initial PDB structures (***Figure 1***) were first cleaned to preserve only the protein part and then solvated with TIP3P water molecules and 0.15 M KCl using CHARMM-GUI [58,59]. The CHARMM36 force field [60] was used, and all simulations were conducted using the GPU-accelerated GROMACS 2019 [61]. The non-bonded forces were calculated using a cut-off of 12 Å and a smoothing switching function to attenuate forces from 10 Å. The particle mesh Ewald (PME) [62] algorithm was employed for long-range electrostatic interactions. All hydrogen-containing bonds were constrained with the LINCS [63]. For equilibration, all systems were first energy minimized for 5000 steps using the steepest descent algorithm and then were subjected to another 125000 steps of restrained NVT (constant particles, constant volume, and constant temperature) simulations with 1 fs as the integration timestep, where the protein backbone and side chains were positionally restrained with harmonic force constants of 400 kJ/mol/Å^2^ and 40 kJ/mol/Å^2^, respectively. The final unrestrained production MD simulations were performed for each system for 200 ns with leap-frog integrator and time step of 2 fs with Nose-Hoover thermostat [64,65] at 303.15 K and Parrinello-Rahman barostat [66] at 1.0 bar and compressibility of 4.5x10^-5^ bar. Snapshots were saved every 100 ps. To calculate the residue RMSF profile, all frames were first aligned with respect to the first frame using backbone atoms and then RMSF values were then calculated for all Cα atoms (***Figure S2***).

### Input Features and Preprocessing

Four types of input features were used for model training: one-hot encoding of amino acids, amino acid index (AAIndex), Rosetta energy terms, and RMSF from MD simulations, as summarized in ***Table S1***. For the graph convolutional network (GCN), the structure of each protein used to initiate MD simulations was used for the construction of the graph. The AAIndex database consists of a set of 566 numerical values describing various physicochemical and biochemical properties for each canonical amino acid [42]. Due to strong correlations among many properties, only 19 principal components (with 100% variance) from PCA analysis of the original AAIndex features[67] were used in the current work after the min-max normalization to the range of [0, 1]. For Rosetta energies (19 components and the total ΔΔG) features, the tanh squashing function [50] was first used to squash energy features greater than 50 REUs (Rosetta energy units), as exceedingly large energy values often arise from steric clashes and are not quantitatively meaningful. Additionally, energy features with small variances were discarded, and only 8 features were used in the final modeling training and testing (***Table S1 and Figure S13***). Finally, the min-max normalization was applied to the selected 7 energy features and the total ΔΔG. The Cα RMSF values from MD simulations were similarly min-max normalized. Lastly, sequence positional information using one-hot encoding was expressed as a binary vector of length 21 (20 for natural amino acid and the extra one for the stop codon).

### ML model design, training and evaluation

We examined the effects of including Rosetta and MD-derived physical features in the four machine learning algorithms previously evaluated in the NN4dms study [41], namely LR, NN, CNN, and GCN. The leaky ReLU activation function was used in NN, CNN, and GCN, and a dropout rate of 0.2 after the dense layer was used to prevent overfitting. In CNN, the strides were [1,1] and the padding was valid. For the implementation, python 3.6 and TensorFlow [68] v1.14 were used. In the GCN, the input graph for GCN was constructed based on the distance matrix of Cα (for glycine) and C_β_ (other residues) with a threshold of 6 Å for Bgl3 and 7 Å for other proteins (see the hyperparameter search below). As illustrated in ***Figure 2***, physical features can be directly added to the input layer, which increases the feature dimension from 40 to 49 per residue(8 Rosetta energies + 1 RMSF value) (***Table S1***). Similarly in the case of GCN, the Rosetta features were added as node features.

**Figure 2.**
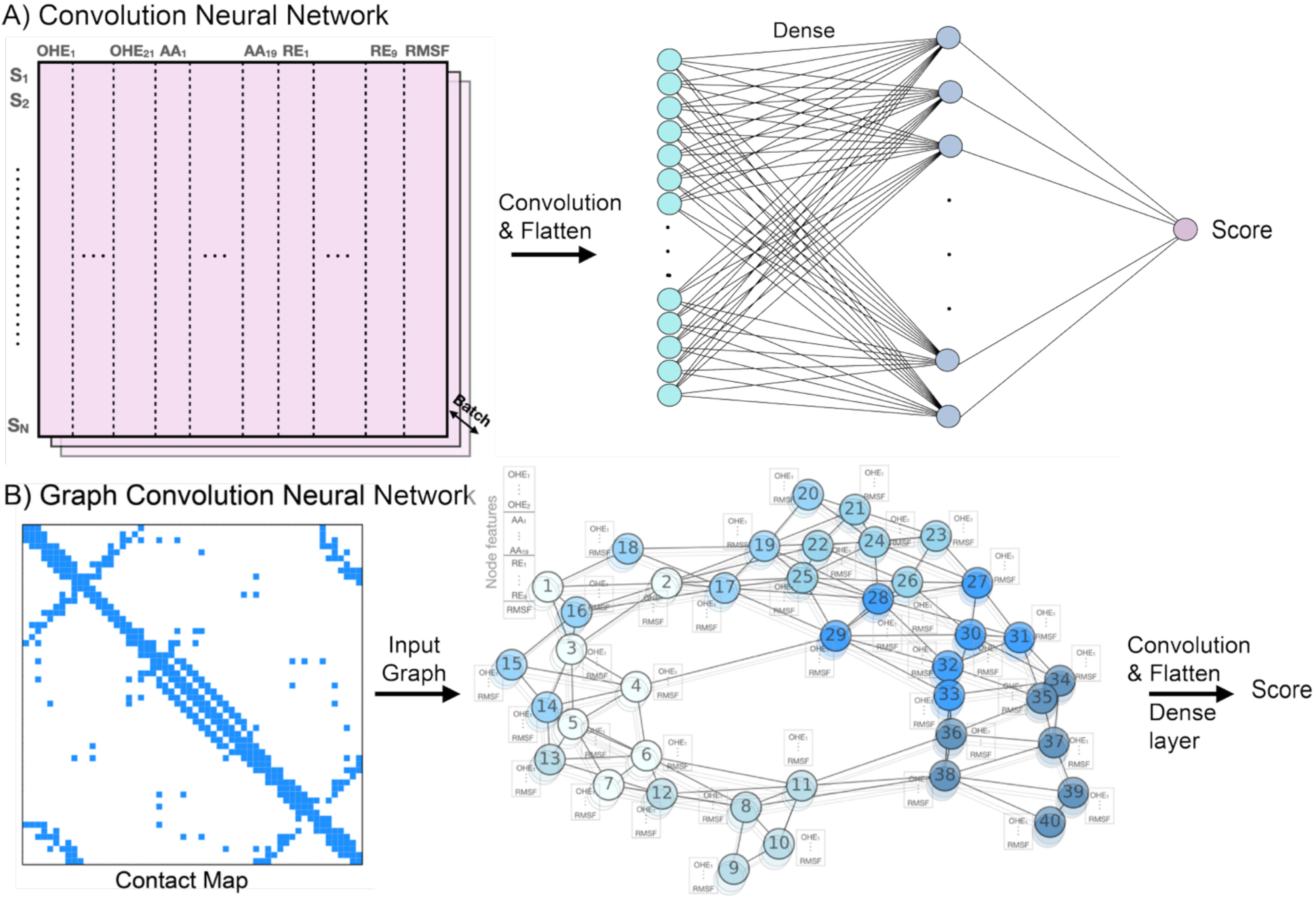
Machine learning architectures for protein variant effect prediction. In CNN, the input dimension is *N*res x 40 without biophysics-based features and *N*res x 49 with biophysics-based features (*N*res: number of residues). In GCN, the wild-type protein structure was used to derive the contact map as graph, where nodes represent residues and edges represent contacts with certain distance cut-off. *Si*: the residue index, OHE: one-hot encoding, AA: amino acid index, RE: Rosetta energies, and RSMF: root mean square fluctuations (from MD).

The random splitting of the dataset was done with 70% for training, 15% for validation, and 15% for test. For studies of dataset size dependence, smaller datsets were created by randomly sampling between 100 datapoints and up to 80% of the full dataset. The remaining 20% was consistently used for validation and testing across all experiments, ensuring that the validation and test sets remained the same for every subset. In mutational extrapolation, all unique mutations from the dataset were collected and randomly divided into training (70%), validation (15%), and testing (15%) sets. All variants containing the corresponding mutations were then assigned to the appropriate set. At last all the overlapped variants between the training, testing, and validation were dropped based on unique mutations in the variant. In positional extrapolation, a similar process was used except that splitting was based on positions.

Hyperparameter search was performed on the validation dataset for all models using grid search with different combination of batch size, learning rate, number of hidden layers, number of nodes, filters and graph threshold. The final selected hyperparameters are reported for the main models are shown in ***Table S2***. Each model was trained five times with different splits with different random seeds. Pearson and Spearman correlation coefficients were calculated using the SciPy library [69] and mean absolute errors (MAE) were calculated using the scikit-learn [70] between the true and predicted scores.

To evaluate how the quality of the DMS dataset affects VEP performance, the resampling experiments were performed in a similar manner as described in the NN4dms study [41]. First, the testing set was constructed from randomly selected 10,000 datapoints from the original GB1 dataset. The left-over dataset was used for construction of randomly resampled libraries for training. For each library, multinomial probability distributions were created for both the input and selected sets by normalizing the original read counts of each variant. These distributions provide the probability of generating a read for any given variant. The fraction of reads was computed in both the sets in the base dataset to decide reads sampled from input and selected sets. The sampling of reads was based on this fraction and Enrich2 scores were calculated for each resampled dataset. These datasets were generated five times for each combination of library size and number of reads from the remaining dataset.

## Results

### Efficient evaluation of mutational effects on protein interactions

The balance between preserving structural integrity and certain degrees of flexibility of a protein is crucial for its function. This intricate balance is governed by tertiary arrangements of residues of different physiochemical properties and their interaction networks. Substitutions of residues can reshape this balance, influence the free energy landscape of protein folding and unfolding, modulate the conformational dynamics, and ultimately affect function. Developing an efficient approach for evaluating the impacts of a given mutation on the physical basis of protein function is thus key to developing robust VEP models [71]. The Rosetta software [49], in particular, can be used to estimate ΔΔG of protein folding stability upon mutation [72] and has been extensively validated using several protein mutation datasets [73]. Importantly, the Rosetta all-atom energy function [49] includes both physics-based and empirical terms to allow quantifications of the mutational effects on key local physical interactions such as electrostatics, hydrogen-bonding, van der Waals interactions, and side chain rotamer states.

Reliable estimation of ΔΔG requires relaxation of the protein conformation in response to mutation, which are generally achieved using either FastRelax or PackRotamerMover protocols in Rosetta. In the FastRelax protocol [56], generally five repeated simulated annealing processes are performed, which consists of re-packing of side chains within a cutoff radius of a selected site followed by global minimization of all dihedral angles while ramping up the repulsive terms in the energy function. However, in the PackRotamerMover protocol [74], only sidechains within the “repacking” radius of the mutated site are allowed to re-pack and no other degrees of freedom are optimized. FastRelax requires longer computing time up to 10s of minutes, depending on the location of the mutation site and the protein size, whereas PackRotamerMover takes only seconds. Importantly, restricted to local side chain rotamer resampling in the PackRotamerMover protocol avoids the challenges of consistently capturing non-trivial global structural relaxation, allowing more reliable description of the energetic consequences of a mutation. Furthermore, the VEPs in this work will be trained with individual Rosetta energy components and local minimization of PackRotamerMover provides a more faithful picture of how the mutation affect various types of local interactions. Of note, the input features include residue RMSFs derived from MD simulations, which inform the ability of different sites of the protein to undergo further (backbone) relaxation to accommodate various mutations (see Methods & ***Figure S2***).

We further optimize the PackRotamerMover protocol to ensure reliable estimation of mutational energetics, focusing on two key parameters: the repacking radius (default: 8 Å) and the number of complete repacking runs before outputting the best energy (default *nloop* = 1). Different repacking radii directly affect the range of residues surrounding the mutated site to be selected and have their side chain rotamers repacked in the protocol, as illustrated in ***Figure 3A-3D***. The results show that a larger repacking radius of 12 Å is necessary to sufficiently optimize the local side chain configurations (***Figures S3-4)*** and obtain reliable ΔΔG estimates (***Figure 3F***; ***Figure S5***). The latter is especially true for mutation to bulkier residues in crowded environments such as T382W (***Figure 3F; Figure S4***). In contrast, the choice of *nloops* of 1, 2 and 3 has no significant impact on the ΔΔG results in all cases examined (***Figure S5***), indicating that the default of 1 iteration of rotamer searching is sufficient. Based on the above analysis, a repacking radius of 12 Å and a *nloop* of 1 were used to calculate Rosetta ΔΔG values for all 20 canonical amino acid substitutions (including the wild-type) at all positions of all five proteins.

**Figure 3.**
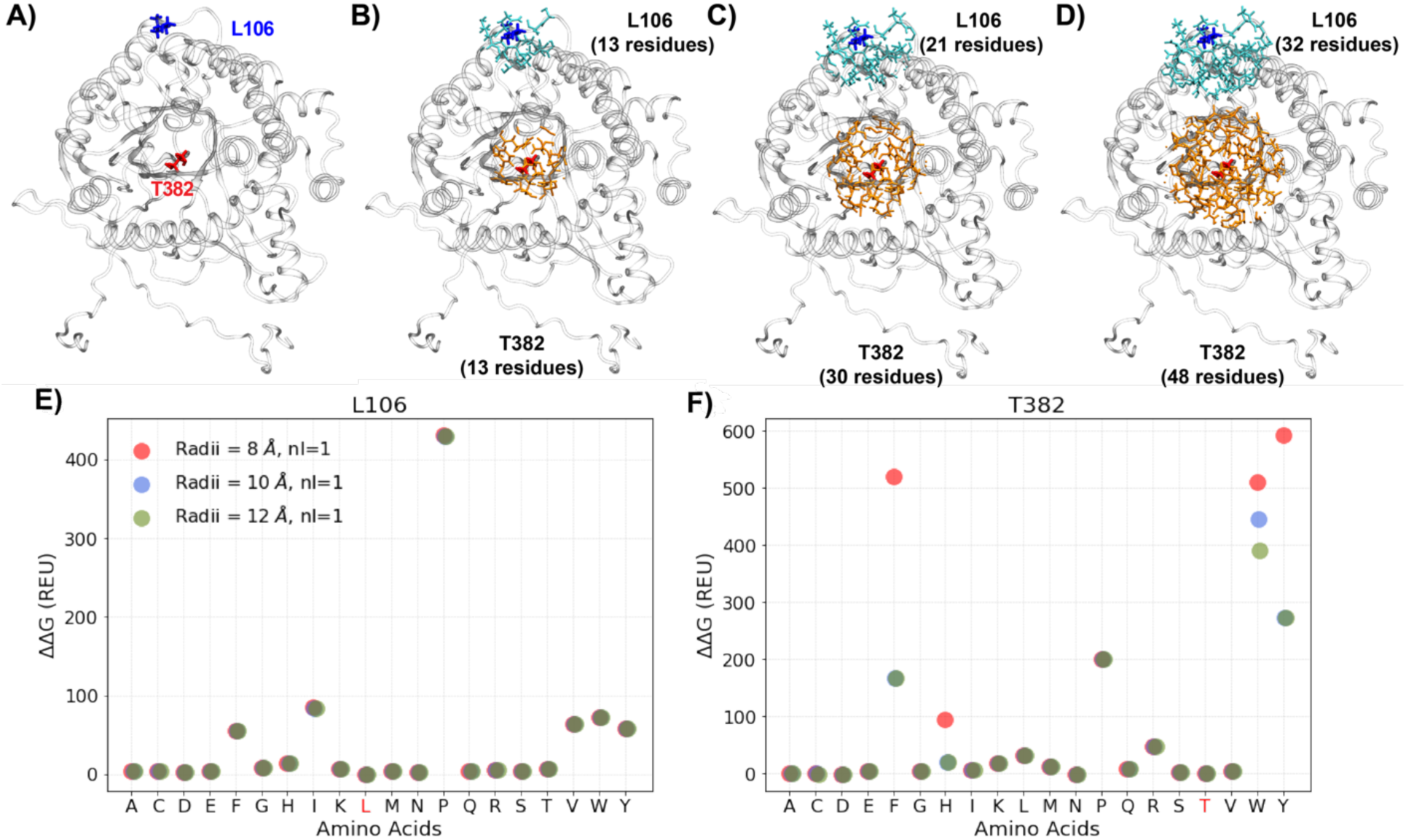
Rosetta local optimization and energy evaluation of site-specific mutations using PackRotamerMover. **A)** The structure of protein Bgl3 in grey cartoon, with a solvent-exposed residue L106 and a buried residue T382 shown in blue and red sticks, respectively. **B-D)** Side chains within 8 Å (B), 10 Å (C), and 12 Å (D) of the two mutation sites, L106 and T382, shown in blue and orange sticks, respectively. The numbers of residues selected by different repacking radii are noted. **E, F)** The ΔΔG values (in REU) for the 20 amino acid substitution at site L106 and T382, calculated using three repacking radii and *nloop* = 1. The wild-type residues are marked in red in the x-axis.

### Effects of biophysics-based features in VEP performance

Armed with the Rosetta energy terms and MD-derived RMSF profiles, we examined the effects of including these biophysics-based features on VEP performance. Models based on LR, NN, CNN, and GCN were trained on the five DMS datasets individually with or without adding the biophysics-based features. All the biophysics-based features are added to the input layer in the with-biophysics models, while the models without biophysics only contain the AAIndex principal components and one-hot encoding for residues in the protein sequence (***Figure 2***). The performance was evaluated on three tasks including the overall prediction (on randomly selected testing datasets), mutational extrapolation as well as the positional extrapolation (see Methods). Five independent repeats for each splitting scheme were performed.

With random splitting, the training and testing splits have similar coverage of mutation types or sites, particularly due to the presence of many higher-order variants in the DMS datasets (***Figure S1A***). For the GB1 and Pab1 datasets with high mutation coverages (98% and 85%, respectively), all four ML models, with or without adding biophysics-based features, achieved nearly perfect Pearson’s correlation (*r* ∼ 0.92 - 0.99) on both training and testing datasets on all five individual splits (***Figure 4*** & ***Figure S6, Table 2***). However, for the Ube4b and Bgl3 datasets with only 63% and 38% mutation coverages (***Figure S1B***), all models were only able to achieve very modest *r* ∼ 0.59 - 0.65 on training and *r* ∼ 0.51 - 0.56 correlations on testing, with only marginal improvements when including biophysics-based features. This modest performance is somewhat surprising for Ube4b given its good 63% mutation coverage. It may be attributed to its poorly packed structure with long loops (∼30% of the sequence) (***Figure 1C***). As a result, Ube4b is highly dynamic (see RMSF in ***Figure S2C***), rendering it difficult to accurately capture mutational impacts on structure and protein energetics. Interestingly, even though the avGFP dataset has only 43% mutation coverage, all models trained on either with or without biophysics-based features achieved generally good correlation (*r* ∼ 0.94 – 0.97) with the LR model showing a slightly lower correlation (*r* ∼ 0.82) (***Figure 4*** & ***Figure S6***). This may be attributed to the tightly packed beta-barrel structure (***Figure 1D***) and the elevated presence of high-order variants (***Figure S1A***). Further analysis of the mean absolute errors (MAEs), however, shows that the addition of biophysics-based terms reduces the MAE for LR and NN models for single mutation variants in protein GB1 and Pab1 datasets (***Figure S9***). Nonetheless, LR and NN frequently underperform compared to CNN and GCN, especially for variants with large numbers of mutations (e.g., ***Figure S9D***). With all five cases considered, including biophysics-based features only demonstrated marginal improvement on the training and testing performance for all models with random splits.

**Figure 4.**
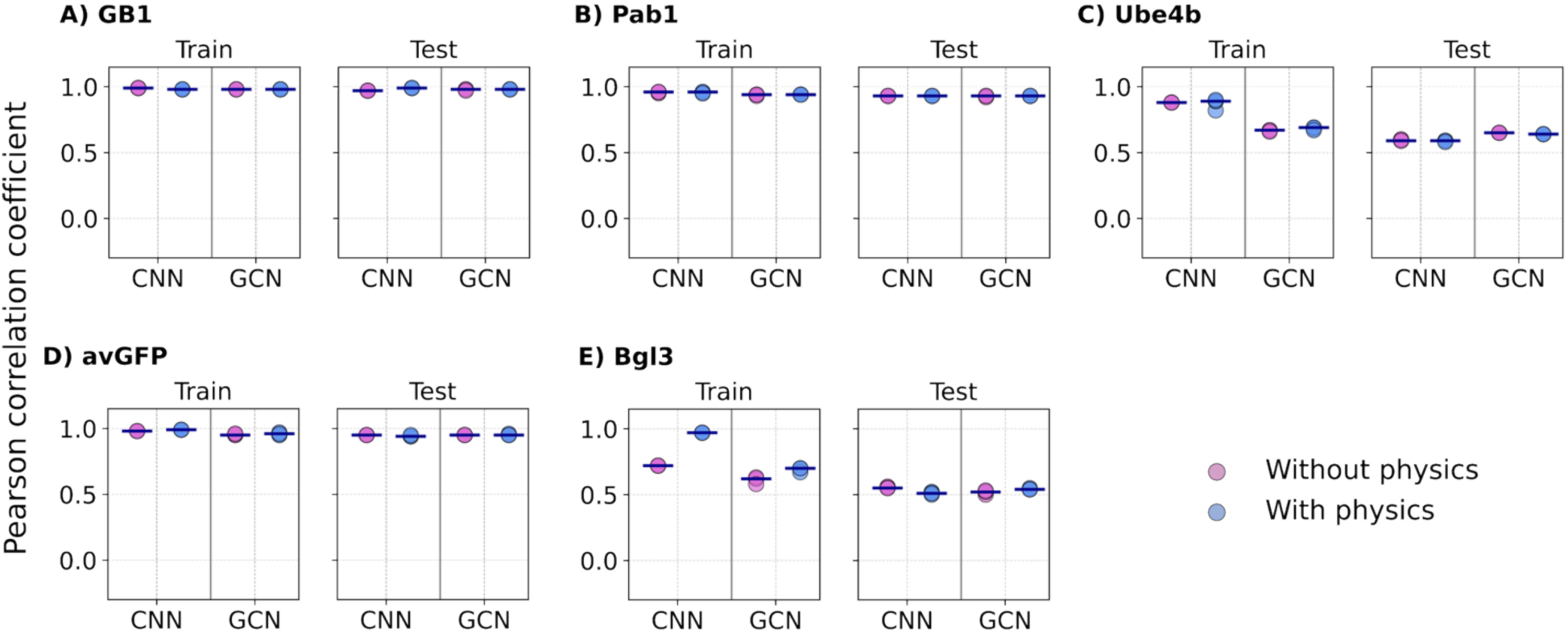
Performance of CNN and GCN models on random split. The 5 proteins are arranged based on decreasing mutational coverage. A) GB1, B) Pab1, C) Ube4b, D) avGFP, and E) Bgl3. Each point indicates one of the 5 random train/test, with the medium shown as black lines.

**Table 2:** Average Pearson correlation coefficients for random, mutational, and positional splits across the four ML models with and without biophysics-based features (BP).

|  |  | Random |  |  |  | Mutational |  |  |  | Positional |  |  |  |
| --- | --- | --- | --- | --- | --- | --- | --- | --- | --- | --- | --- | --- | --- |
|  |  | LR | NN | CNN | GCN | LR | NN | CNN | GCN | LR | NN | CNN | GCN |
| GB1 | No BP | 0.92 | 0.97 | 0.97 | 0.97 | 0.68 | 0.80 | 0.89 | 0.90 | -0.01 | -0.02 | 0.05 | 0.37 |
|  | BP | 0.92 | 0.98 | 0.99 | 0.97 | 0.80 | 0.85 | 0.89 | 0.90 | -0.17 | -0.07 | **0.43** | 0.40 |
| Pab1 | No BP | 0.90 | 0.92 | 0.93 | 0.93 | 0.44 | 0.67 | 0.76 | 0.71 | 0.01 | 0.08 | 0.17 | 0.03 |
|  | BP | 0.91 | 0.94 | 0.93 | 0.93 | 0.69 | 0.70 | 0.76 | 0.74 | 0.10 | 0.08 | **0.48** | **0.57** |
| Ube4b | No BP | 0.60 | 0.65 | 0.59 | 0.65 | 0.19 | 0.31 | 0.45 | 0.37 | -0.03 | 0.02 | 0.08 | 0.10 |
|  | BP | 0.60 | 0.65 | 0.59 | 0.64 | **0.40** | 0.43 | 0.46 | 0.44 | -0.00 | -0.04 | 0.21 | 0.25 |
| avGFP | No BP | 0.82 | 0.96 | 0.94 | 0.95 | 0.06 | 0.28 | 0.64 | 0.52 | -0.01 | -0.06 | 0.15 | 0.07 |
|  | BP | 0.82 | 0.97 | 0.94 | 0.95 | **0.61** | **0.64** | 0.79 | 0.74 | 0.08 | 0.28 | **0.69** | **0.63** |
| Bgl3 | No BP | 0.52 | 0.49 | 0.55 | 0.52 | 0.02 | 0.10 | 0.18 | 0.30 | 0.00 | 0.01 | 0.11 | 0.10 |
|  | BP | 0.56 | 0.56 | 0.51 | 0.54 | **0.33** | 0.33 | 0.33 | 0.34 | 0.00 | -0.02 | 0.31 | 0.22 |

### Biophysics-based features improve VEP performance against small datasets

To further evaluate model robustness against the training set size, we trained series of CNN and GCN models with and without biophysics-based features for all proteins using only 100 randomly selected datapoints up to 0.9 fraction of the total datasets. The results, summarized in ***Figure S12***, clearly show that models trained with biophysics-based features generally perform better than the without-biophysics counterparts when only seeing the limited training dataset size, especially when the sizes are less than 1000 (with low to moderate *r*). It is interesting to note that, in the case of Ube4b, CNN outperformed GCN. This suggests that if the protein structure contain many flexible loops, including structural information provides little to no advantage to the model performance, especially when trained with limited number of data points.

Besides the DMS dataset size and mutation coverage, the quality of fitness or functional scores can also impact VEP training and performance. The fitness scores were mainly calculated using read frequencies from sequencing of the initial library and the isolated variants after the functional selection. There is a tradeoff between the dataset size and average reads per variant when the budget for the total sequencing number is fixed. We designed resampling experiments of the most complete DMS dataset of GB1 following the same protocol outlined in the NN4DMS study [41] and evaluated the performance of CNN and GCN models trained with biophysics-based features. The resample experiment generated DMS subsets with varying library sizes and numbers of reads from the original dataset [54,52]. As summarized in ***Figure 5***, both small library sizes (thus not enough mutation coverage) and small numbers of reads (thus unreliable fitness scores) can dramatically diminish the accuracy of trained VEPs. However, including biophysics-based features clearly allow both CNN and GCN models to perform more robustly as the quality of the DMS dataset deteriorates due to either small library size or DNA reads per variant (lower left corners in ***Figure 5***). Compared to the best CNN models previous reported in the NN4DMS study [41] and the biophysics-based CNN, when number of sequencing reads are low almost every protein library size has been increased modestly for biophysics-based CNN. For other larger sequencing reads performance of both the models are similar overall. Similarly in the case of the biophysics-based GCN, showed moderate performance improvements for low sequencing read counts and across protein libraries; however, with protein library size 5e4 and 1e5, performance decreased slightly. Curiously, both CNN and GCN architecture appears to perform similar overall. Taken together, for random splits where the training and testing data have high overlaps, including biophysics-based features marginally affects VEP performance with large DMS datasets, but can substantially improve the robustness of trained models when the mutational coverage is limited and/or the number of DNA sequencing reads is small.

**Figure 5.**
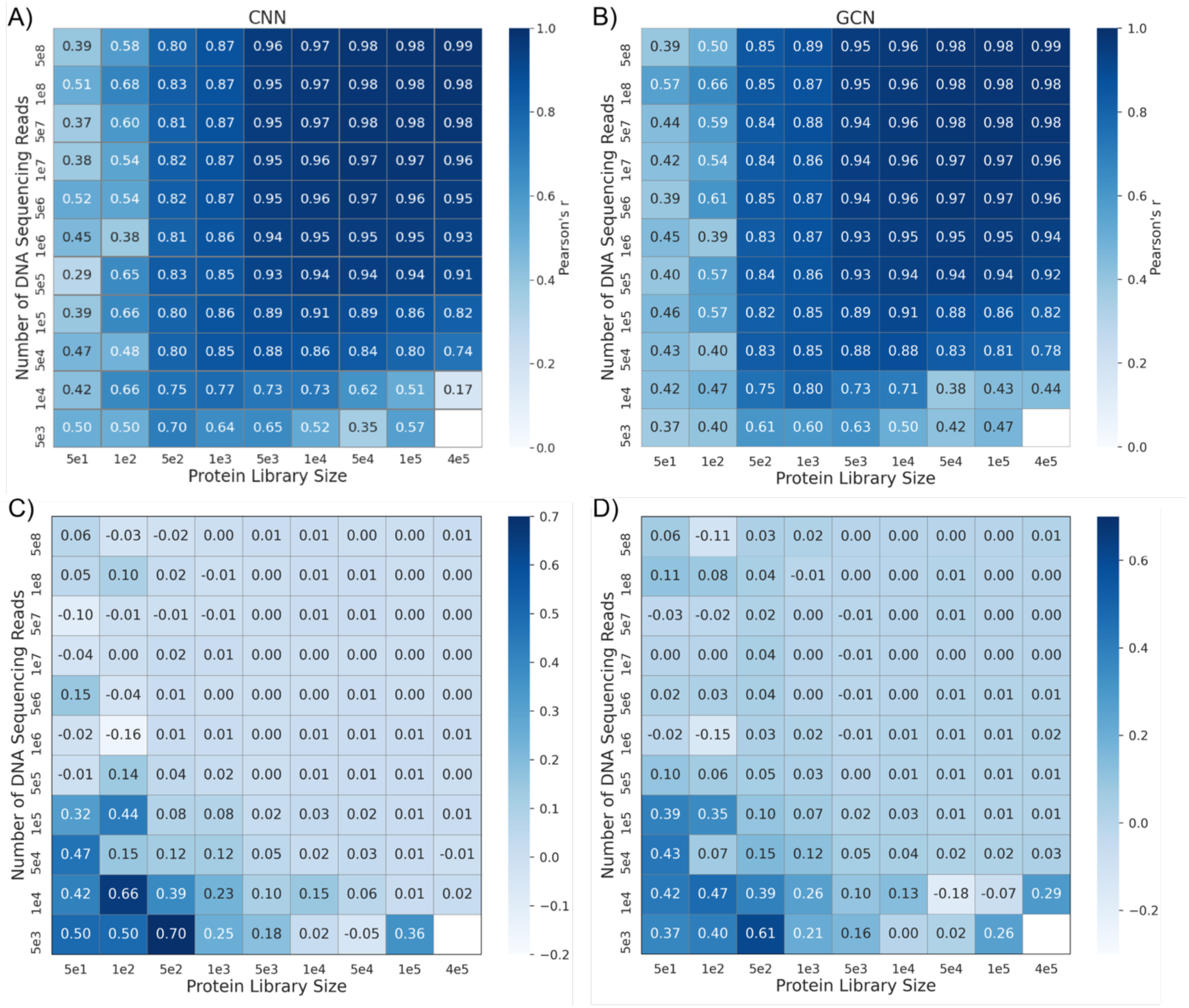
Dependence of VEP performance on the library size and number of sequencing reads. A, B) Pearson’s correlations for CNN and GCN models trained with biophysics-based features using different combination of the library size and number of sequencing reads for the protein GB1 DMS dataset. The empty box represents the combination where the number of variants were not enough for the experiment. C, D) Differences in Pearson’s correlations between the above CNN and GCN models and previously CNN models trained without biophysics-based features [41].

### Biophysics significantly enhances mutational and positional extrapolation

A major challenge identified from the previous NN4DMS study [41] is to reliably predict the effects of novel mutation types or mutation positions not seen in training. In the mutational extrapolation task, a VEP model trained on several mutation types on specific sites is asked to make predictions for other mutation types on those sites (see Methods). As summarized in ***Figure 6, Figure S7*** and ***Table 2***, all models are still performing well with the GB1 dataset due to its 98% sequence percentage converge. For other protein datasets, adding biophysics-based features in general improved the performance of all four ML models on their testing sets, especially for avGFP. For models with lower complexity such as LR and NN, biophysics-based features can dramatically improve prediction, such as LR models for the Pab1 (from 0.44 to 0.69), Ube4b (from 0.19 to 0.40), avGFP (from 0.06 to 0.61) and Bgl3 (from 0.02 to 0.33) datasets and NN models in the avGFP (from 0.28 to 0.64) and Bgl3 (from 0.1 to 0.33) datasets (***Figure S7***). For models with higher complexities such as CNN and GCN, those biophysics-based features still could help achieve slightly better performances, such as in avGFP (from 0.52 to 0.74), and Bgl3 (from 0.18 to 0.33) datasets (***Figure 6***). These results indicate that biophysics-based features provide important and useful information for mutational extrapolation task.

**Figure 6.**
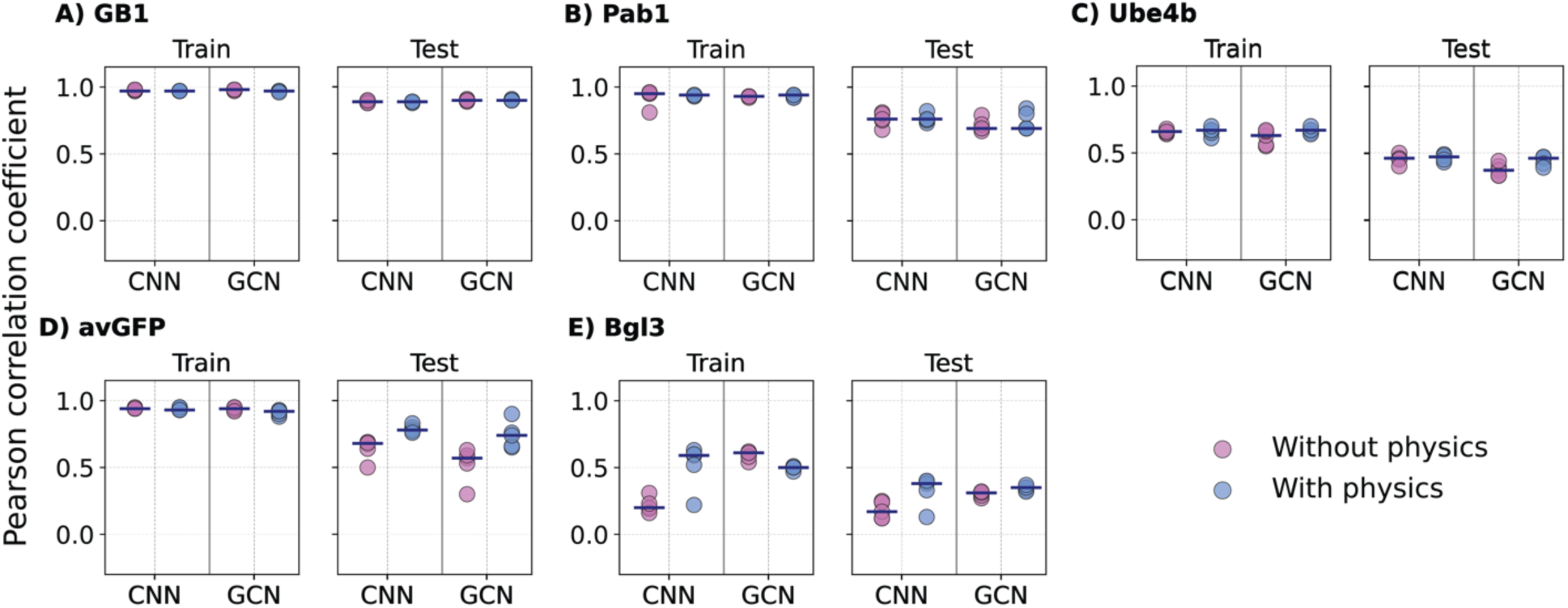
Performance of CNN and GCN models trained with and without biophysics for mutational extrapolation. The 5 proteins are arranged with decreasing mutational coverage. A) GB1, B) Pab1, C) Ube4b, D) avGFP, and E) Bgl3. Each point indicates one of the 5 random mutational train/test, with the medium shown as black lines.

Positional extrapolation is the most challenging task since it requires a VEP model trained only on certain specific mutation sites to make correct predictions on mutations to sites it has never seen during training. All four ML models, when trained without biophysics-based features, show either no correlation (r ∼ 0) or very low correlation (r < 0.2) on the testing set, except for GCN in the case of GB1 (***Figure 7*** & ***Figure S8***), highlighting the difficulty of this task and also the merit of incorporating protein structure information. Strikingly, incorporating biophysics-based features significantly enhances the ability of all VEP models to handle positional extrapolation, especially with CNN and GCN (***Figure 7***). For example, in the case of GB1 CNN (from 0.05 to 0.43), Pab1 (0.17 to 0.48), avGFP (0.15 to 0.69) dataset, and GCN in the Pab1 (0.03 to 0.57) and avGFP (0.07 to 0.63) datasets. In other cases where the protein itself has more complexity to model, adding biophysics-based features can still marginally improve their performance, such as CNN and GCN in the Ube4b (from 0.08-0.10 to 0.21-0.25) and Bgl3 (from 0.1 to 0.22-0.31) dataset. This strongly supports that biophysics-based features are indeed beneficial for positional extrapolation, especially for well-structured proteins with low sequence percentage coverages (such as avGFP). The MAE comparison also shows that incorporating biophysics-based terms reduces the MAE across all models and for all types of variants as shown in ***Figure S9***, ***Figure S10***, and ***Figure S11***.

**Figure 7.**
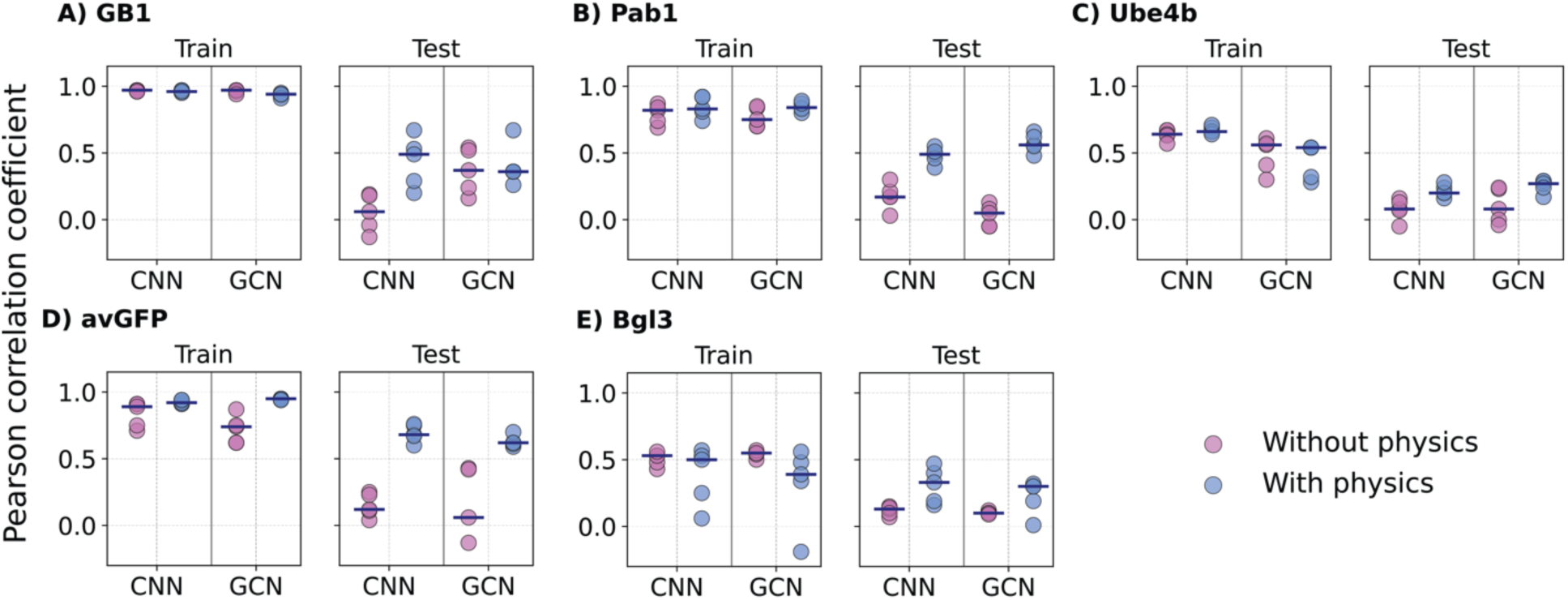
Performance of CNN and GCN models trained with and without biophysics for positional extrapolation. The 5 proteins are arranged with decreasing mutational coverage. A) GB1, B) Pab1, C) Ube4b, D) avGFP, and E) Bgl3. Each point indicates one of the 5 random positional train/test, with the medium shown as black lines

Comparing performance of our simpler biophysics-based model with another biophysics-informed transformer model, the local METL model [48], we observe that in the case of mutational extrapolation, our model achieves a Spearman correlation of 0.62 for GFP (METL: ∼0.7), 0.43 for Ube4b (METL: 0.5), ∼0.9 for GB1 (METL: ∼0.9), and ∼0.75 for Pab1 (METL: 0.8). For positional extrapolation, our model attains a correlation of only 0.45 compared to METL which achieved ∼0.7 for GB1, 0.5 for GFP (METL: ∼0.58), 0.56 for Pab1 (METL: ∼0.59), and 0.26 for Ube4b (METL: ∼0.2). Overall, our model achieves comparable performance to the local METL model in most cases except for GB1 positional extrapolation, suggesting that overall incorporating biophysics-based features can maintain performance levels similar to those of more complex transformer-based approaches.

## Discussion

Characterization of protein sequence-to-function relationships is critical to study protein evolution, understand genetic diseases, and design proteins with enhanced or new functions. However, the inherent complexity from large sequence space, structural variability, and conformational dynamics makes this mapping task extremely challenging. The availability of sequencing data, protein structures, and high-throughput functional assays has allowed the training of machine learning models to infer the protein sequence-function relationships and predict variation effects. In this work, we develop and evaluate a simple and efficient approach for incorporating biophysics-based features to train VEP models with superior robustness against dataset limitations and ability to predict effects of novel mutational types or positions. This approach only requires the number of residues x 19 fast Rosetta structural relaxation and energy evaluations, each of which takes only seconds for a typical protein. The effectiveness of the biophysics-inspired VEP development approach has been evaluated using existing DMS datasets of five proteins with diverse size, topology and mutation coverage. The results strongly support the effectiveness of incorporating biophysics to improve VEP performance and overcome DMS dataset limitations, especially for positional extrapolation with both CNN and GCN.

Our results provide several key insights for building robust VEP models trained on DMS datasets. Firstly, the sequence percentage coverage is critical. For the GB1 dataset, which has 98% coverage, all models, including simple LR performed well in the random and mutational splitting schemes. Whereas for the Bgl3 datasets, which has only 38% coverage, all VEP models showed medium to low correlation, with average of ∼0.54 for random and ∼0.33 for mutational. Arguably, such data scarcity could only be effectively mitigated in general by acquiring more functionally labelled data points. Secondly, incorporating the physical principles of protein structure and interaction is an effective strategy for overcoming data scarcity problem in VEP development. The physical principles could be captured by estimating mutational impacts on the protein energetics and by considering the structural flexibility as derived from short MD simulations. Models trained with these biophysics-based features can dramatically enhance mutational and positional extrapolations for well-structured proteins, such as for the avGFP dataset with only 43% mutation coverage. Last but not the least, the structure and dynamics of the protein are also important factors that affect VEP performance. For example, existing variants may not effectively inform on new ones for highly dynamic proteins, such as in the case of Ube4b. In these cases, obtaining reliable biophysics-based features is also much more challenging. The current approach focuses on capturing localized structural and energetic impacts of mutations changes would only help marginally. Exploring the use of advanced machine learning techniques, such as transformer-based models, in combination with biophysics-based features, could further improve extrapolation capabilities.

In conclusion, our study demonstrates that incorporating biophysics-based features into deep learning models significantly enhances their ability to predict protein function from sequence data, especially in extrapolation scenarios. This approach holds great promise for advancing protein engineering and understanding protein-related diseases. These predictors can help as valuable tools for researchers, aiding the prioritization and comprehensive understanding of the genetic variations for various diseases and protein engineering. It can also contribute to identifying potential drug targets and understanding them under various conditions. It can help predict the impact of sequence modifications on protein behavior and functionality.

## Acknowledgements

The authors thank Sam Gelman and Anthony Gitter for help discussions. This work is supported by NIH R35 GM144045 (to Chen).

## Author contributions

Barethiya, Guan, and Chen conceived the study. Barethiya, Huang and Liu performed modeling and analysis. Barethiya, Huang and Chen wrote the manuscript with inputs from Liu and Guan.

## Competing interests

The authors declare no competing interests.

## Supplementary information

All training data and scripts are available on GitHub (https://github.com/SBarethiya/VEP). The online version contains supplementary material available at https://doi.org/xxxx

## Notes

### Competing Interest Statement

The authors have declared no competing interest.

### Summary of Updates

The revision includes the supplementary materials that was not originally uploaded by the journal.

## Reference

1. Maheshri N, Koerber JT, Kaspar BK, Schaffer DV. Directed evolution of adeno-associated virus yields enhanced gene delivery vectors. Nat Biotechnol. 2006;24: 198–204. doi:10.1038/nbt1182

2. Findlay GM, Daza RM, Martin B, Zhang MD, Leith AP, Gasperini M, et al. Accurate classification of BRCA1 variants with saturation genome editing. Nature. 2018;562: 217–222. doi:10.1038/s41586-018-0461-z

3. Lee JM, Huddleston J, Doud MB, Hooper KA, Wu NC, Bedford T, et al. Deep mutational scanning of hemagglutinin helps predict evolutionary fates of human H3N2 influenza variants. Proceedings of the National Academy of Sciences. 2018;115: E8276–E8285. doi:10.1073/pnas.1806133115

4. Campbell RE, Tour O, Palmer AE, Steinbach PA, Baird GS, Zacharias DA, et al. A monomeric red fluorescent protein. Proceedings of the National Academy of Sciences. 2002;99: 7877– 7882. doi:10.1073/pnas.082243699

5. Mahlich Y, Steinegger M, Rost B, Bromberg Y. HFSP: high speed homology-driven function annotation of proteins. Bioinformatics. 2018;34: i304–i312. doi:10.1093/bioinformatics/bty262

6. Loewenstein Y, Raimondo D, Redfern OC, Watson J, Frishman D, Linial M, et al. Protein function annotation by homology-based inference. Genome Biology. 2009;10: 207. doi:10.1186/gb-2009-10-2-207

7. Kankainen M, Ojala T, Holm L. BLANNOTATOR: enhanced homology-based function prediction of bacterial proteins. BMC Bioinformatics. 2012;13: 33. doi:10.1186/1471-2105-13-33

8. Fowler DM, Araya CL, Fleishman SJ, Kellogg EH, Stephany JJ, Baker D, et al. High-resolution mapping of protein sequence-function relationships. Nat Methods. 2010;7: 741–746. doi:10.1038/nmeth.1492

9. Fowler DM, Fields S. Deep mutational scanning: a new style of protein science. Nat Methods. 2014;11: 801–807. doi:10.1038/nmeth.3027

10. Fowler DM, Stephany JJ, Fields S. Measuring the activity of protein variants on a large scale using deep mutational scanning. Nat Protoc. 2014;9: 2267–2284. doi:10.1038/nprot.2014.153

11. Romero PA, Tran TM, Abate AR. Dissecting enzyme function with microfluidic-based deep mutational scanning. Proceedings of the National Academy of Sciences. 2015;112: 7159– 7164. doi:10.1073/pnas.1422285112

12. Kitzman JO, Starita LM, Lo RS, Fields S, Shendure J. Massively parallel single-amino-acid mutagenesis. Nat Methods. 2015;12: 203–206. doi:10.1038/nmeth.3223

13. Melamed D, Young DL, Gamble CE, Miller CR, Fields S. Deep mutational scanning of an RRM domain of the Saccharomyces cerevisiae poly(A)-binding protein. RNA. 2013;19: 1537–1551. doi:10.1261/rna.040709.113

14. Weile J, Sun S, Cote AG, Knapp J, Verby M, Mellor JC, et al. A framework for exhaustively mapping functional missense variants. Molecular Systems Biology. 2017;13: 957. doi:10.15252/msb.20177908

15. Mighell TL, Evans-Dutson S, O’Roak BJ. A Saturation Mutagenesis Approach to Understanding PTEN Lipid Phosphatase Activity and Genotype-Phenotype Relationships. The American Journal of Human Genetics. 2018;102: 943–955. doi:10.1016/j.ajhg.2018.03.018

16. Zhang L, Sarangi V, Ho M-F, Moon I, Kalari KR, Wang L, et al. *SLCO1B1*: Application and Limitations of Deep Mutational Scanning for Genomic Missense Variant Function. Drug Metabolism and Disposition. 2021;49: 395–404. doi:10.1124/dmd.120.000264

17. Wei H, Li X. Deep mutational scanning: A versatile tool in systematically mapping genotypes to phenotypes. Front Genet. 2023;14. doi:10.3389/fgene.2023.1087267

18. Durbin RM, Altshuler D, Durbin RM, Abecasis GR, Bentley DR, Chakravarti A, et al. A map of human genome variation from population-scale sequencing. Nature. 2010;467: 1061–1073. doi:10.1038/nature09534

19. Berman HM, Westbrook J, Feng Z, Gilliland G, Bhat TN, Weissig H, et al. The Protein Data Bank. Nucleic Acids Res. 2000;28: 235–242. doi:10.1093/nar/28.1.235

20. Jumper J, Evans R, Pritzel A, Green T, Figurnov M, Ronneberger O, et al. Highly accurate protein structure prediction with AlphaFold. Nature. 2021;596: 583–589. doi:10.1038/s41586-021-03819-2

21. Baek M, DiMaio F, Anishchenko I, Dauparas J, Ovchinnikov S, Lee GR, et al. Accurate prediction of protein structures and interactions using a three-track neural network. Science. 2021;373: 871–876. doi:10.1126/science.abj8754

22. Lin Z, Akin H, Rao R, Hie B, Zhu Z, Lu W, et al. Evolutionary-scale prediction of atomic-level protein structure with a language model. Science. 2023;379: 1123–1130. doi:10.1126/science.ade2574

23. Ioannidis NM, Rothstein JH, Pejaver V, Middha S, McDonnell SK, Baheti S, et al. REVEL: An Ensemble Method for Predicting the Pathogenicity of Rare Missense Variants. The American Journal of Human Genetics. 2016;99: 877–885. doi:10.1016/j.ajhg.2016.08.016

24. Hopf TA, Ingraham JB, Poelwijk FJ, Schärfe CPI, Springer M, Sander C, et al. Mutation effects predicted from sequence co-variation. Nat Biotechnol. 2017;35: 128–135. doi:10.1038/nbt.3769

25. Gray VE, Hause RJ, Luebeck J, Shendure J, Fowler DM. Quantitative Missense Variant Effect Prediction Using Large-Scale Mutagenesis Data. Cell Systems. 2018;6: 116–124.e3. doi:10.1016/j.cels.2017.11.003

26. Xu Y, Verma D, Sheridan RP, Liaw A, Ma J, Marshall NM, et al. Deep Dive into Machine Learning Models for Protein Engineering. J Chem Inf Model. 2020;60: 2773–2790. doi:10.1021/acs.jcim.0c00073

27. Horne J, Shukla D. Recent Advances in Machine Learning Variant Effect Prediction Tools for Protein Engineering. Ind Eng Chem Res. 2022;61: 6235–6245. doi:10.1021/acs.iecr.1c04943

28. Dunham AS, Beltrao P, AlQuraishi M. High-throughput deep learning variant effect prediction with Sequence UNET. Genome Biology. 2023;24: 110. doi:10.1186/s13059-023-02948-3

29. Freschlin CR, Fahlberg SA, Heinzelman P, Romero PA. Neural network extrapolation to distant regions of the protein fitness landscape. Nat Commun. 2024;15: 6405. doi:10.1038/s41467-024-50712-3

30. Listov D, Goverde CA, Correia BE, Fleishman SJ. Opportunities and challenges in design and optimization of protein function. Nat Rev Mol Cell Biol. 2024;25: 639–653. doi:10.1038/s41580-024-00718-y

31. Hartman EC, Tullman-Ercek D. Learning from protein fitness landscapes: a review of mutability, epistasis, and evolution. Current Opinion in Systems Biology. 2019;14: 25–31. doi:10.1016/j.coisb.2019.02.006

32. Chen L, Zhang Z, Li Z, Li R, Huo R, Chen L, et al. Learning protein fitness landscapes with deep mutational scanning data from multiple sources. Cell Systems. 2023;14: 706–721.e5. doi:10.1016/j.cels.2023.07.003

33. Johnson MS, Reddy G, Desai MM. Epistasis and evolution: recent advances and an outlook for prediction. BMC Biology. 2023;21: 120. doi:10.1186/s12915-023-01585-3

34. Shin H, Cho B-K. Rational Protein Engineering Guided by Deep Mutational Scanning. International Journal of Molecular Sciences. 2015;16: 23094–23110. doi:10.3390/ijms160923094

35. Dewachter L, Brooks AN, Noon K, Cialek C, Clark-ElSayed A, Schalck T, et al. Deep mutational scanning of essential bacterial proteins can guide antibiotic development. Nat Commun. 2023;14: 241. doi:10.1038/s41467-023-35940-3

36. Hanning KR, Minot M, Warrender AK, Kelton W, Reddy ST. Deep mutational scanning for therapeutic antibody engineering. Trends in Pharmacological Sciences. 2022;43: 123–135. doi:10.1016/j.tips.2021.11.010

37. Riesselman AJ, Ingraham JB, Marks DS. Deep generative models of genetic variation capture the effects of mutations. Nat Methods. 2018;15: 816–822. doi:10.1038/s41592-018-0138-4

38. Shin J-E, Riesselman AJ, Kollasch AW, McMahon C, Simon E, Sander C, et al. Protein design and variant prediction using autoregressive generative models. Nat Commun. 2021;12: 2403. doi:10.1038/s41467-021-22732-w

39. Song H, Bremer BJ, Hinds EC, Raskutti G, Romero PA. Inferring Protein Sequence-Function Relationships with Large-Scale Positive-Unlabeled Learning. Cell Systems. 2021;12: 92–101.e8. doi:10.1016/j.cels.2020.10.007

40. Luo Y, Jiang G, Yu T, Liu Y, Vo L, Ding H, et al. ECNet is an evolutionary context-integrated deep learning framework for protein engineering. Nat Commun. 2021;12: 5743. doi:10.1038/s41467-021-25976-8

41. Gelman S, Fahlberg SA, Heinzelman P, Romero PA, Gitter A. Neural networks to learn protein sequence–function relationships from deep mutational scanning data. Proceedings of the National Academy of Sciences. 2021;118: e2104878118. doi:10.1073/pnas.2104878118

42. Kawashima S, Pokarowski P, Pokarowska M, Kolinski A, Katayama T, Kanehisa M. AAindex: amino acid index database, progress report 2008. Nucleic Acids Research. 2008;36: D202– D205. doi:10.1093/nar/gkm998

43. Mater AC, Sandhu M, Jackson C. The NK Landscape as a Versatile Benchmark for Machine Learning Driven Protein Engineering. bioRxiv; 2020. p. 2020.09.30.319780. doi:10.1101/2020.09.30.319780

44. Rao RM, Liu J, Verkuil R, Meier J, Canny J, Abbeel P, et al. MSA Transformer. Proceedings of the 38th International Conference on Machine Learning. PMLR; 2021. pp. 8844–8856. Available: https://proceedings.mlr.press/v139/rao21a.html

45. Yang Y, Perdikaris P. Physics-informed deep generative models. arXiv; 2018. doi:10.48550/arXiv.1812.03511

46. Raissi M, Perdikaris P, Karniadakis GE. Physics-informed neural networks: A deep learning framework for solving forward and inverse problems involving nonlinear partial differential equations. Journal of Computational Physics. 2019;378: 686–707. doi:10.1016/j.jcp.2018.10.045

47. Zheng L-E, Barethiya S, Nordquist E, Chen J. Machine Learning Generation of Dynamic Protein Conformational Ensembles. Molecules. 2023;28: 4047. doi:10.3390/molecules28104047

48. Gelman S, Johnson B, Freschlin C, D’Costa S, Gitter A, Romero PA. Biophysics-based protein language models for protein engineering. 2024. doi:10.1101/2024.03.15.585128

49. Alford RF, Leaver-Fay A, Jeliazkov JR, O’Meara MJ, DiMaio FP, Park H, et al. The Rosetta All-Atom Energy Function for Macromolecular Modeling and Design. J Chem Theory Comput. 2017;13: 3031–3048. doi:10.1021/acs.jctc.7b00125

50. Nordquist E, Zhang G, Barethiya S, Ji N, White KM, Han L, et al. Incorporating physics to overcome data scarcity in predictive modeling of protein function: A case study of BK channels. PLOS Computational Biology. 2023;19: e1011460. doi:10.1371/journal.pcbi.1011460

51. Sarkisyan KS, Bolotin DA, Meer MV, Usmanova DR, Mishin AS, Sharonov GV, et al. Local fitness landscape of the green fluorescent protein. Nature. 2016;533: 397–401. doi:10.1038/nature17995

52. Olson CA, Wu NC, Sun R. A Comprehensive Biophysical Description of Pairwise Epistasis throughout an Entire Protein Domain. Current Biology. 2014;24: 2643–2651. doi:10.1016/j.cub.2014.09.072

53. Starita LM, Pruneda JN, Lo RS, Fowler DM, Kim HJ, Hiatt JB, et al. Activity-enhancing mutations in an E3 ubiquitin ligase identified by high-throughput mutagenesis. Proceedings of the National Academy of Sciences. 2013;110: E1263–E1272. doi:10.1073/pnas.1303309110

54. Fowler DM, Araya CL, Gerard W, Fields S. Enrich: software for analysis of protein function by enrichment and depletion of variants. Bioinformatics. 2011;27: 3430–3431. doi:10.1093/bioinformatics/btr577

55. Romero PA, Tran TM, Abate AR. Dissecting enzyme function with microfluidic-based deep mutational scanning. Proceedings of the National Academy of Sciences. 2015;112: 7159– 7164. doi:10.1073/pnas.1422285112

56. Tyka MD, Keedy DA, André I, DiMaio F, Song Y, Richardson DC, et al. Alternate States of Proteins Revealed by Detailed Energy Landscape Mapping. Journal of Molecular Biology. 2011;405: 607–618. doi:10.1016/j.jmb.2010.11.008

57. Chaudhury S, Lyskov S, Gray JJ. PyRosetta: a script-based interface for implementing molecular modeling algorithms using Rosetta. Bioinformatics. 2010;26: 689–691. doi:10.1093/bioinformatics/btq007

58. Jo S, Kim T, Iyer VG, Im W. CHARMM-GUI: A web-based graphical user interface for CHARMM. Journal of Computational Chemistry. 2008;29: 1859–1865. doi:10.1002/jcc.20945

59. Lee J, Cheng X, Swails JM, Yeom MS, Eastman PK, Lemkul JA, et al. CHARMM-GUI Input Generator for NAMD, GROMACS, AMBER, OpenMM, and CHARMM/OpenMM Simulations Using the CHARMM36 Additive Force Field. J Chem Theory Comput. 2016;12: 405–413. doi:10.1021/acs.jctc.5b00935

60. Huang J, MacKerell Jr AD. CHARMM36 all-atom additive protein force field: Validation based on comparison to NMR data. Journal of Computational Chemistry. 2013;34: 2135–2145. doi:10.1002/jcc.23354

61. Abraham MJ, Murtola T, Schulz R, Páll S, Smith JC, Hess B, et al. GROMACS: High performance molecular simulations through multi-level parallelism from laptops to supercomputers. SoftwareX. 2015;1–2: 19–25. doi:10.1016/j.softx.2015.06.001

62. Darden T, York D, Pedersen L. Particle mesh Ewald: An N⋅log(N) method for Ewald sums in large systems. The Journal of Chemical Physics. 1993;98: 10089–10092. doi:10.1063/1.464397

63. Hess B. P-LINCS: A Parallel Linear Constraint Solver for Molecular Simulation. J Chem Theory Comput. 2008;4: 116–122. doi:10.1021/ct700200b

64. Nosé S, Klein M l. Constant pressure molecular dynamics for molecular systems. Molecular Physics. 1983;50: 1055–1076. doi:10.1080/00268978300102851

65. Hoover WG. Canonical dynamics: Equilibrium phase-space distributions. Phys Rev A. 1985;31: 1695–1697. doi:10.1103/PhysRevA.31.1695

66. Parrinello M, Rahman A. Polymorphic transitions in single crystals: A new molecular dynamics method. Journal of Applied Physics. 1981;52: 7182–7190. doi:10.1063/1.328693

67. Rubin AF, Gelman H, Lucas N, Bajjalieh SM, Papenfuss AT, Speed TP, et al. A statistical framework for analyzing deep mutational scanning data. Genome Biology. 2017;18: 150. doi:10.1186/s13059-017-1272-5

68. Abadi M, Agarwal A, Barham P, Brevdo E, Chen Z, Citro C, et al. TensorFlow: Large-Scale Machine Learning on Heterogeneous Distributed Systems. In: arXiv.org [Internet]. 14 Mar 2016 [cited 9 June 2024]. Available: https://arxiv.org/abs/1603.04467v2

69. Virtanen P, Gommers R, Oliphant TE, Haberland M, Reddy T, Cournapeau D, et al. SciPy 1.0: fundamental algorithms for scientific computing in Python. Nat Methods. 2020;17: 261–272. doi:10.1038/s41592-019-0686-2

70. Pedregosa F, Varoquaux G, Gramfort A, Michel V, Thirion B, Grisel O, et al. Scikit-learn: Machine Learning in Python. arXiv; 2018. doi:10.48550/arXiv.1201.0490

71. Marabotti A, Scafuri B, Facchiano A. Predicting the stability of mutant proteins by computational approaches: an overview. Briefings in Bioinformatics. 2021;22: bbaa074. doi:10.1093/bib/bbaa074

72. Kellogg EH, Leaver-Fay A, Baker D. Role of conformational sampling in computing mutation-induced changes in protein structure and stability. Proteins: Structure, Function, and Bioinformatics. 2011;79: 830–838. doi:10.1002/prot.22921

73. Frenz B, Lewis SM, King I, DiMaio F, Park H, Song Y. Prediction of Protein Mutational Free Energy: Benchmark and Sampling Improvements Increase Classification Accuracy. Front Bioeng Biotechnol. 2020;8. doi:10.3389/fbioe.2020.558247

74. Fleishman SJ, Leaver-Fay A, Corn JE, Strauch E-M, Khare SD, Koga N, et al. RosettaScripts: A Scripting Language Interface to the Rosetta Macromolecular Modeling Suite. PLOS ONE. 2011;6: e20161. doi:10.1371/journal.pone.0020161

