## Supplemental tables and figures for "Overcoming Extrapolation Challenges of Deep Learning by Incorporating Physics in Protein Sequence-Function Modeling"

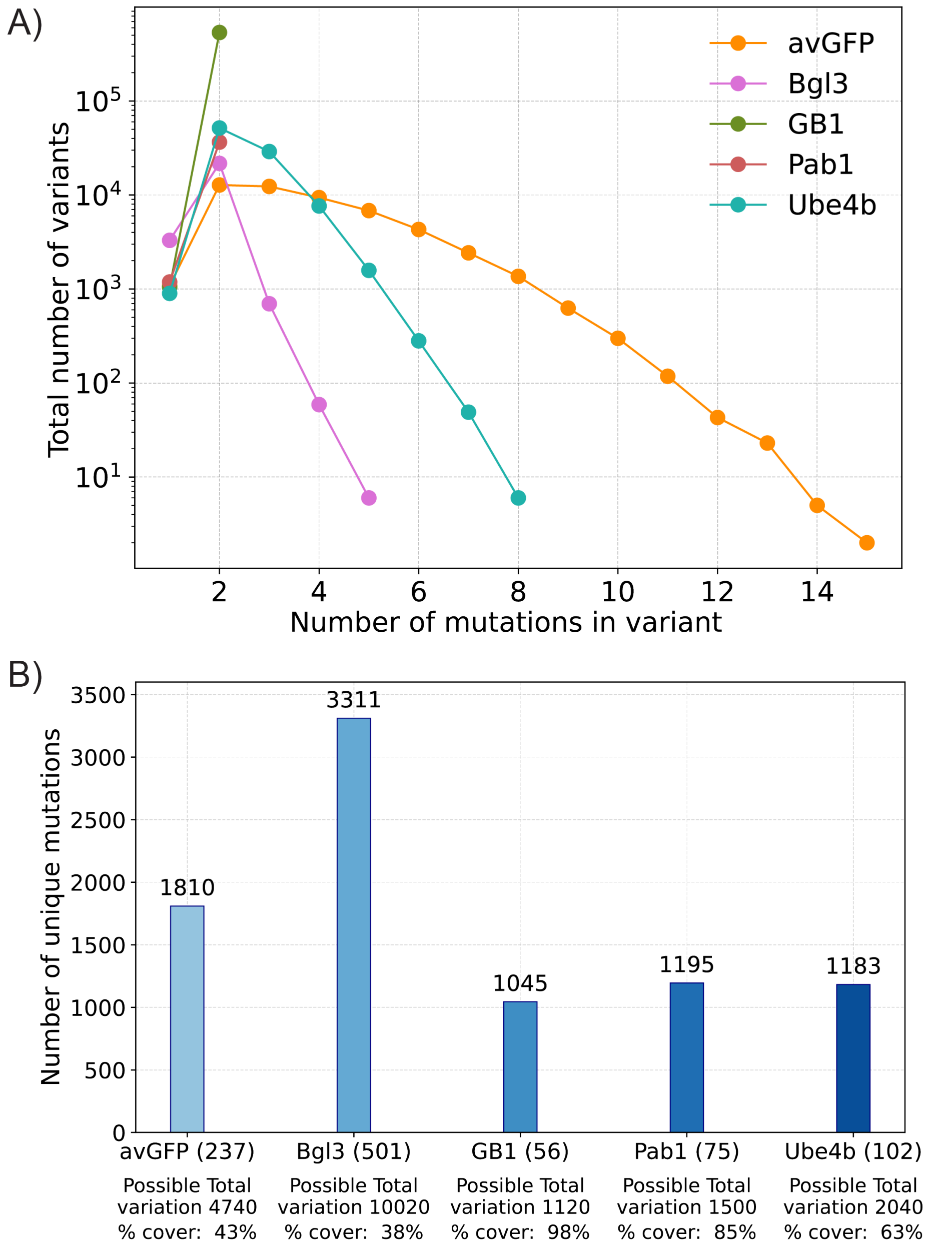


**Figure S1. Distribution and coverage of variants in DMS datasets.** A) Distribution of various containing different number of mutations in each of the five datasets. B) Number of unique mutations sampled for each protein. The coverage percentage, listed under the x-axis, is defined as the number of unique mutations (+ wildtype) divided by the total possible number (20 x sequence length). The sequence length of each protein is given in parathesis next to the protein name.

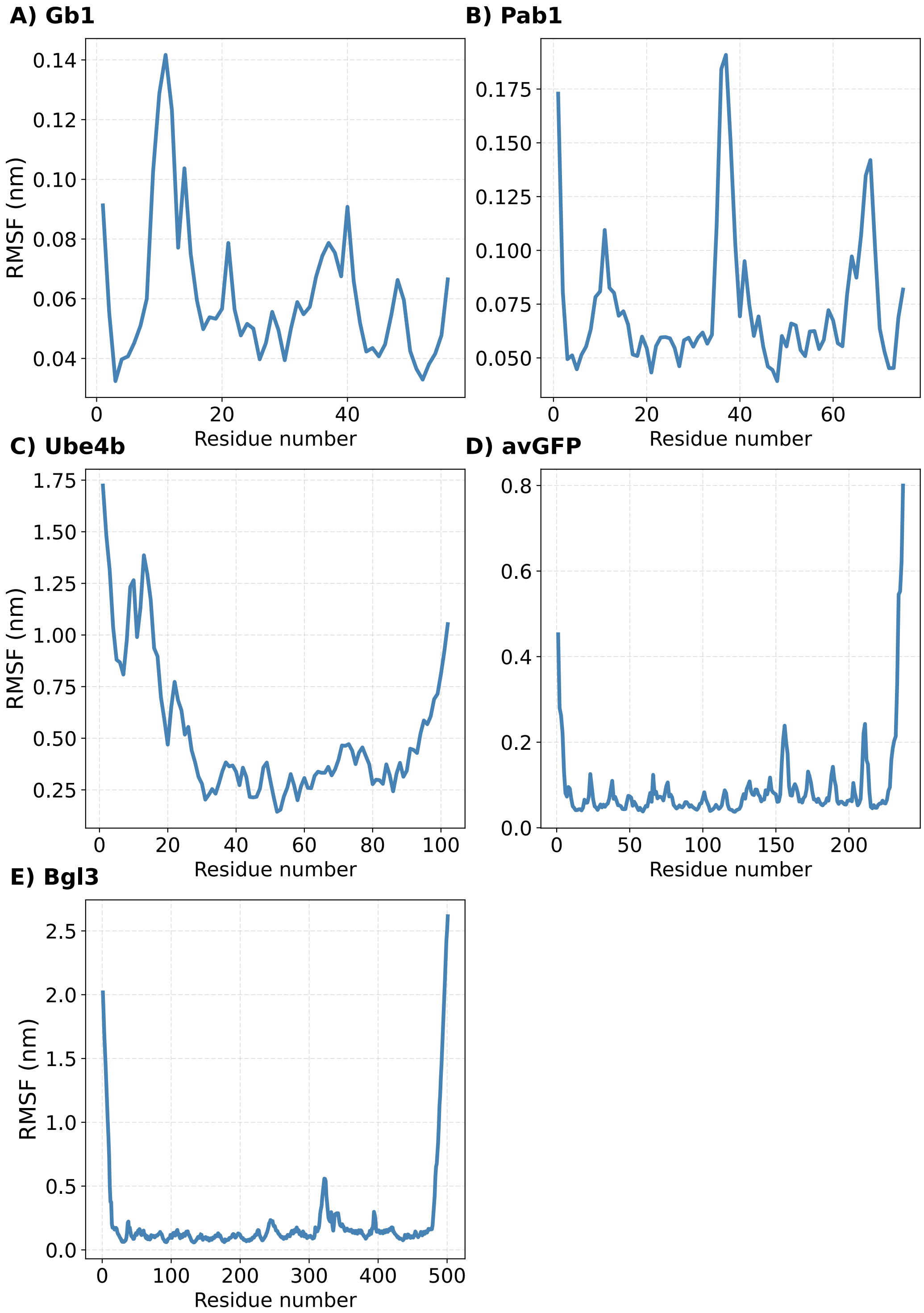


**Figure S2. Residue Root Mean Square Fluctuations (RMSF) profiles of all five proteins,** derived from 200 ns atomistic simulations (see Methods).

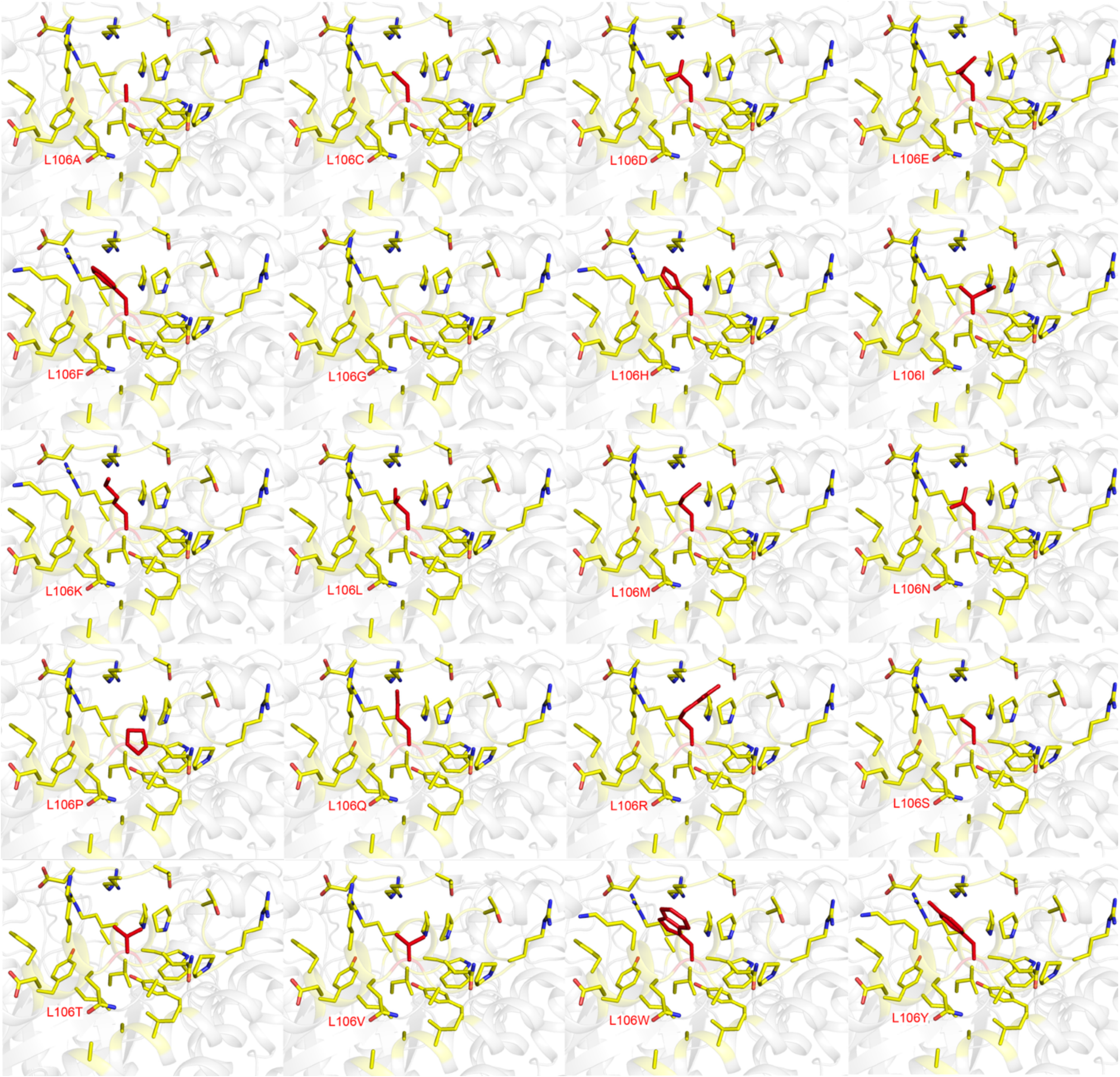


**Figure S3. Optimized local side chain configurations upon mutating solvent exposed L106 of protein Bgl3 to all 19 other amino acids**, generated using the PackRotamerMover protocol with a repacking radius of 12 Å and *nloops* of 1. The protein backbone is shown in white cartoon and all residues within the repacking radius are represented as yellow sticks with residue L106 highlighted in red. Elements are colored as follows: C (yellow), N (blue), O (red), and S (orange).

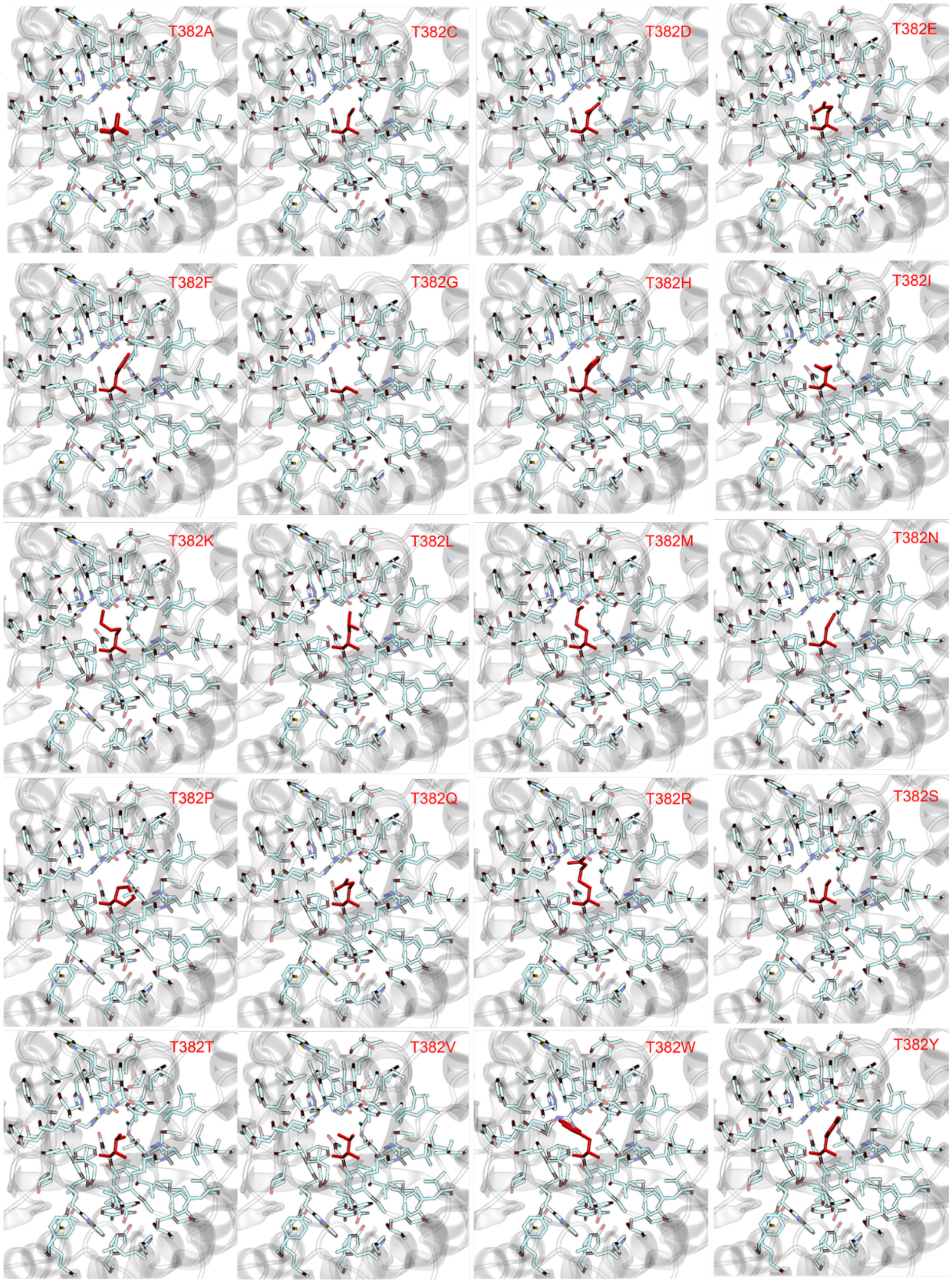


**Figure S4. Optimized local side chain configurations upon mutating buried T382 of protein Bgl3 to all 19 other amino acids**, generated using the PackRotamerMover protocol with a repacking radius of 12 Å and *nloops* of 1. The protein backbone is shown in white cartoon and all residues within the repacking radius are represented as cyan sticks with T382 highlighted in red. Elements are colored as follows: C (cyan), N (blue), O (red), and S (yellow).

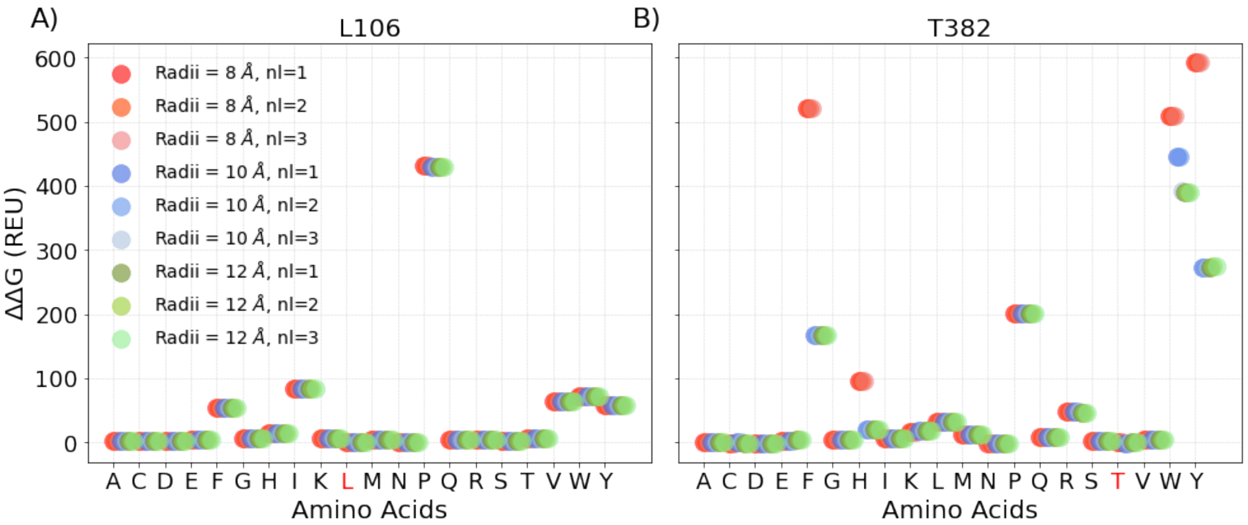


**Figure S5. Convergence of ΔΔG with different repacking radii and nloop values in PackRotamerMover**. The ΔΔG (in REU) values for the 20 amino acid substitution was calculated for solvent exposed L106 (A) and buried T382 (B).

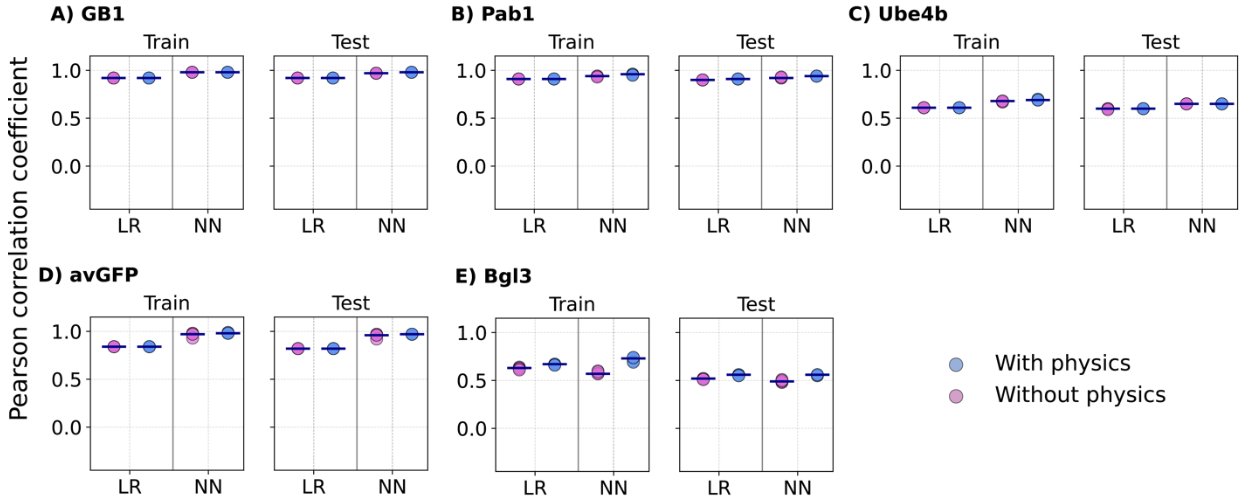


**Figure S6. Performance of LR and NN models with or without biophysics in the random splitting scheme**. The 5 proteins are arranged from the largest to smallest sequence space coverage: A) GB1, B) Pab1, C) Ube4b, D) avGFP, and E) Bgl3. Each point indicates one of the 5 random train/test and the medium of the 5 replicates are marked using black lines.

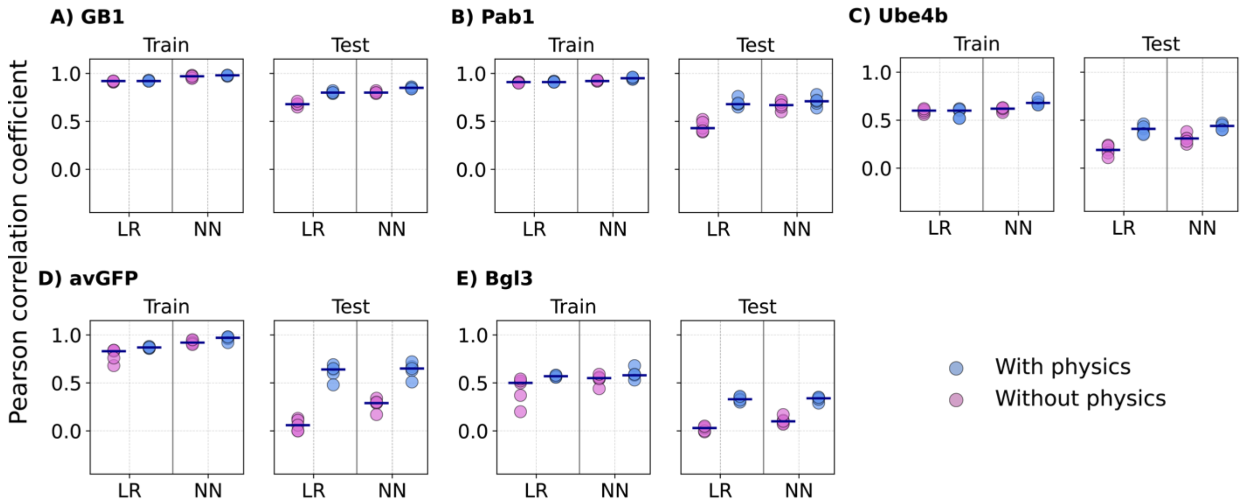


**Figure S7. Performance of LR and NN models with and without biophysics models in the mutational splitting scheme**. The 5 proteins are arranged from the largest to smallest sequence space coverage: A) GB1, B) Pab1, C) Ube4b, D) avGFP, and E) Bgl3. Each point indicates one of the 5 random train/test and the medium of the 5 replicates are marked using black lines.

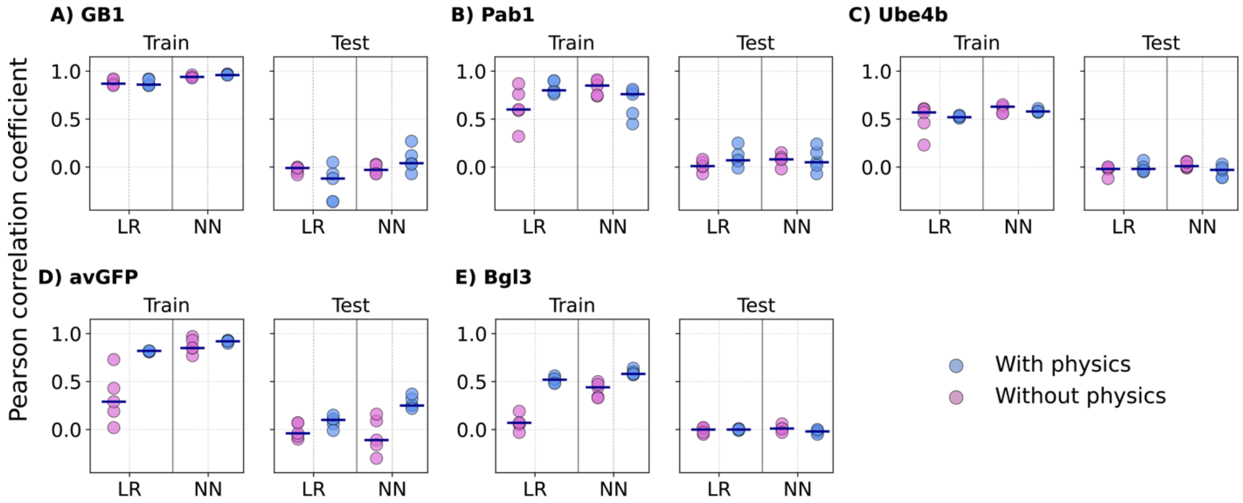


**Figure S8. Performance of LR and NN models with and without biophysics models in the positional splitting scheme**. The 5 proteins are arranged from the largest to smallest sequence space coverage. A) GB1, B) Ube4b, C) avGFP, D) Pab1, and E) Bgl3. Each point indicates one of the 5 random train/test and the medium of the 5 replicates are marked using black lines.

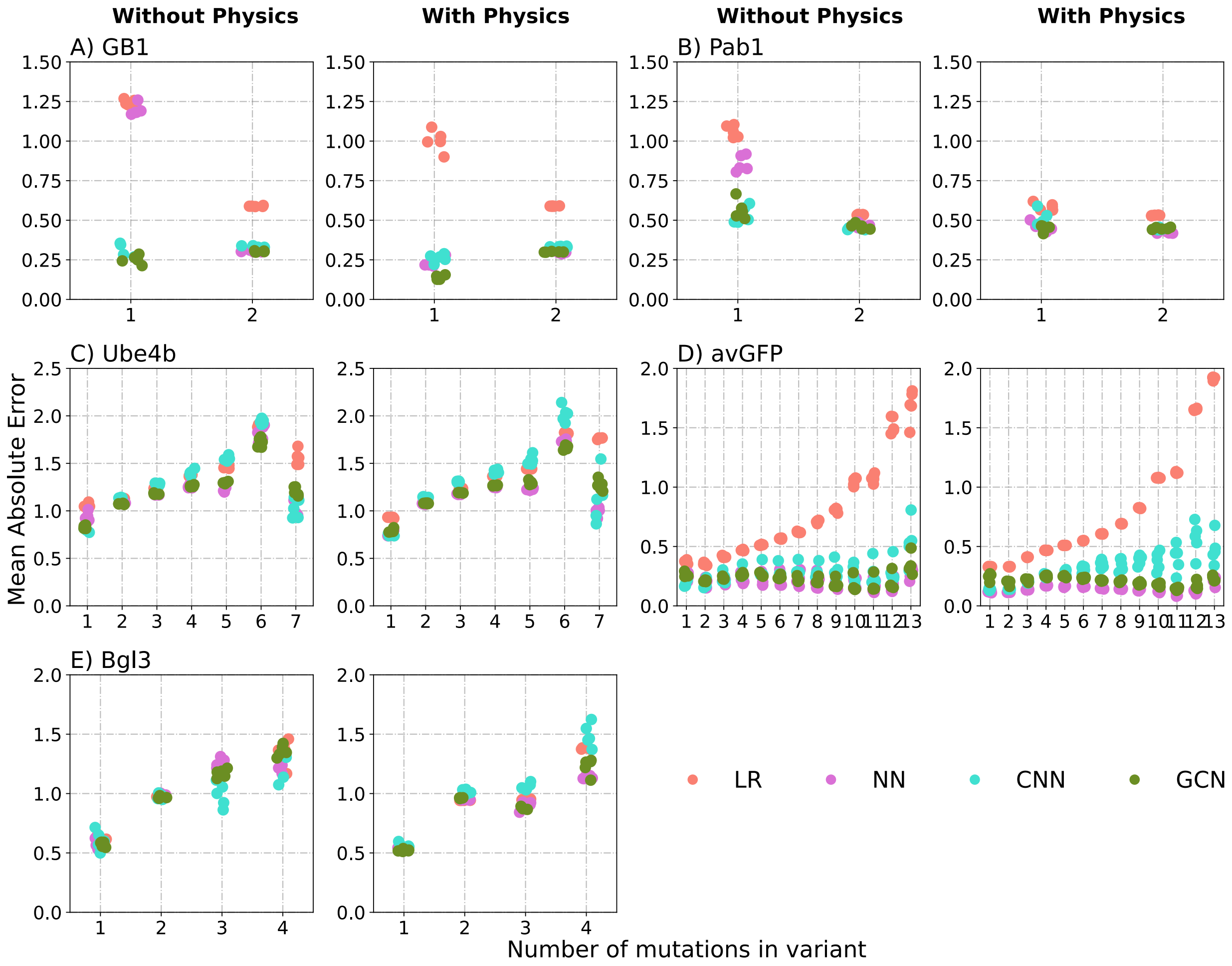


**Figure S9. Mean absolute error with respect to number of mutations in variant for models trained with and without biophysics using random splitting.** Each color point represents one of the 5 random train/test for LR (salmon), NN (orchid), CNN (turquoise) and GCN (green) models.

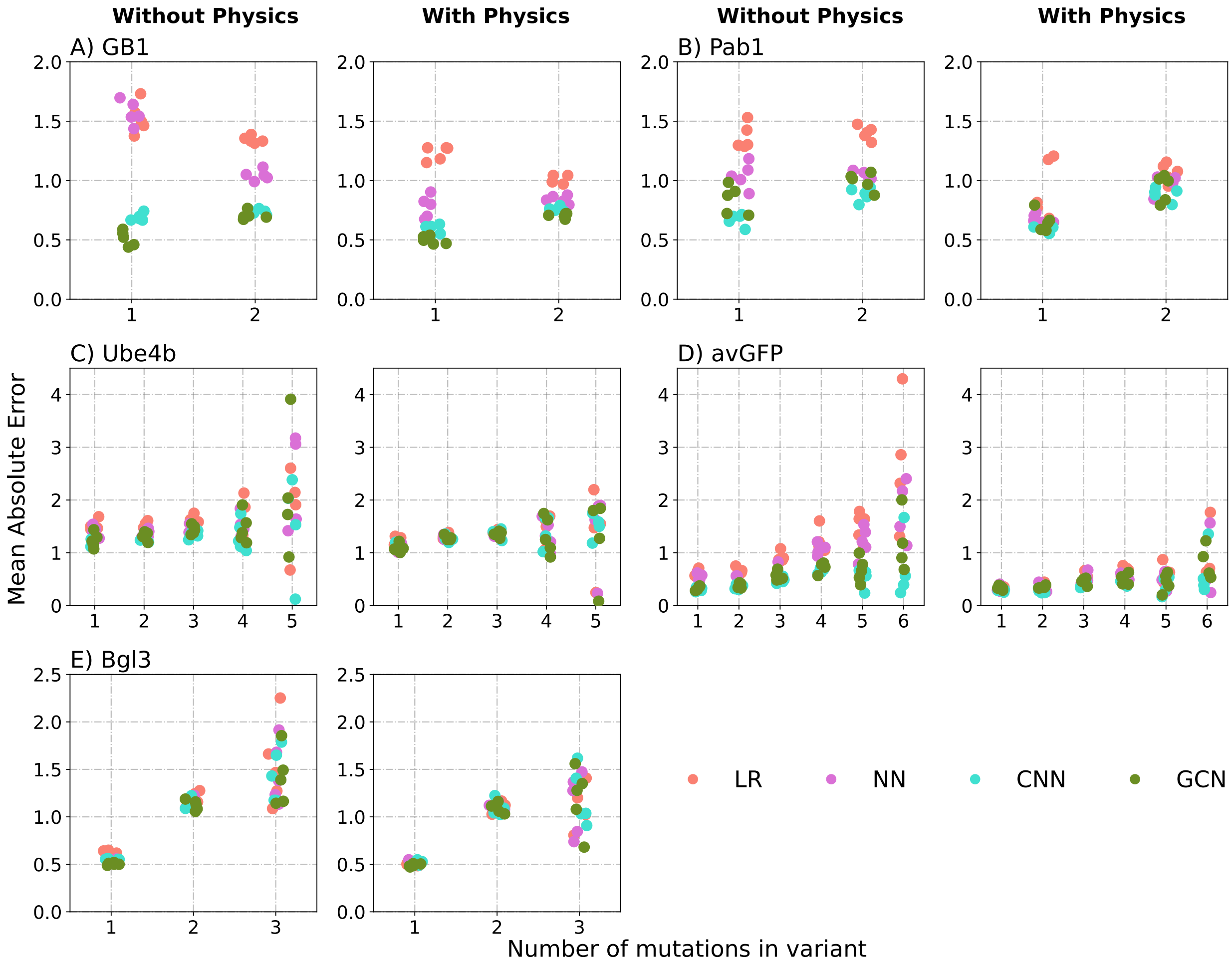


**Figure S10. Mean absolute error with respect to number of mutations in variant for models trained with and without biophysics for mutational extrapolation**. Each color point represents one of the 5 random splits for LR (salmon), NN (orchid), CNN (turquoise) and GCN (green) model.

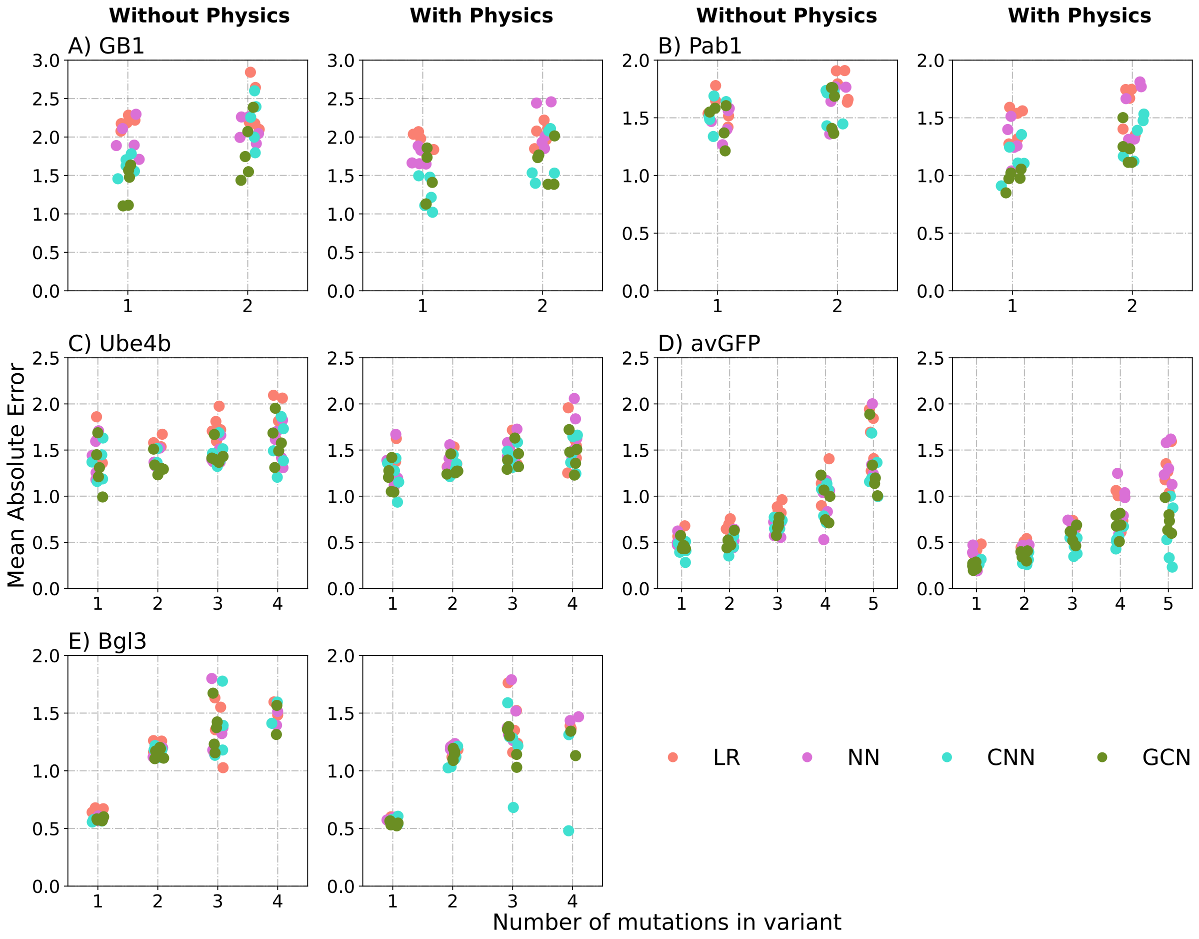


**Figure S11. Mean absolute error with respect to number of mutations in variant for models trained with and without biophysics using random splitting for positional extrapolation**. Each color point represents one of the 5 random splits for LR (salmon), NN (orchid), CNN (turquoise) and GCN (green) model.

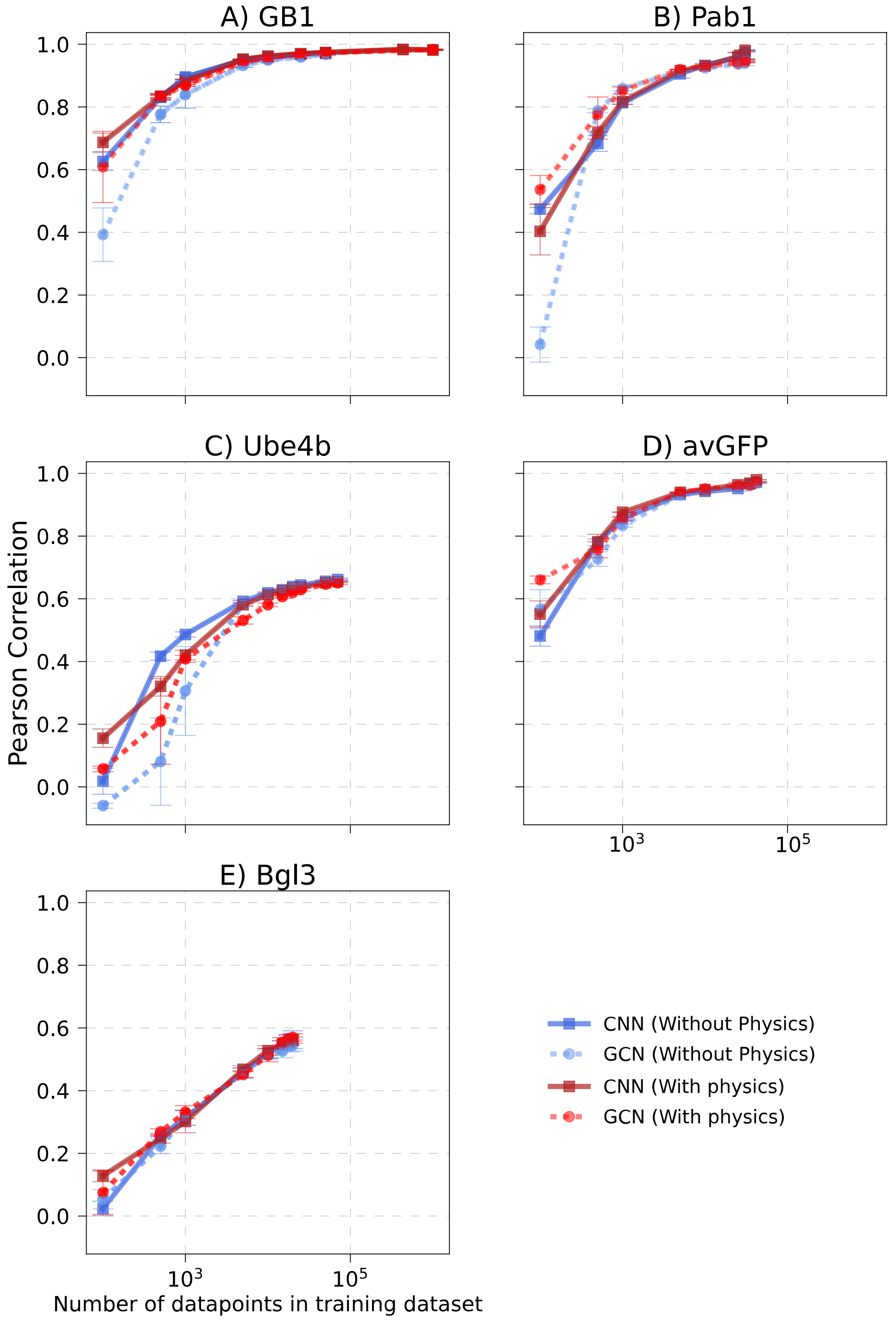


**Figure S12. Performance of CNN and GCN models with and without biophysical features with respect to training random split fraction**. The training dataset was increased from 100 datapoints to 0.9 fraction of the total dataset of each protein. The solid line indicates CNN, and dashed line indicates GCN model.

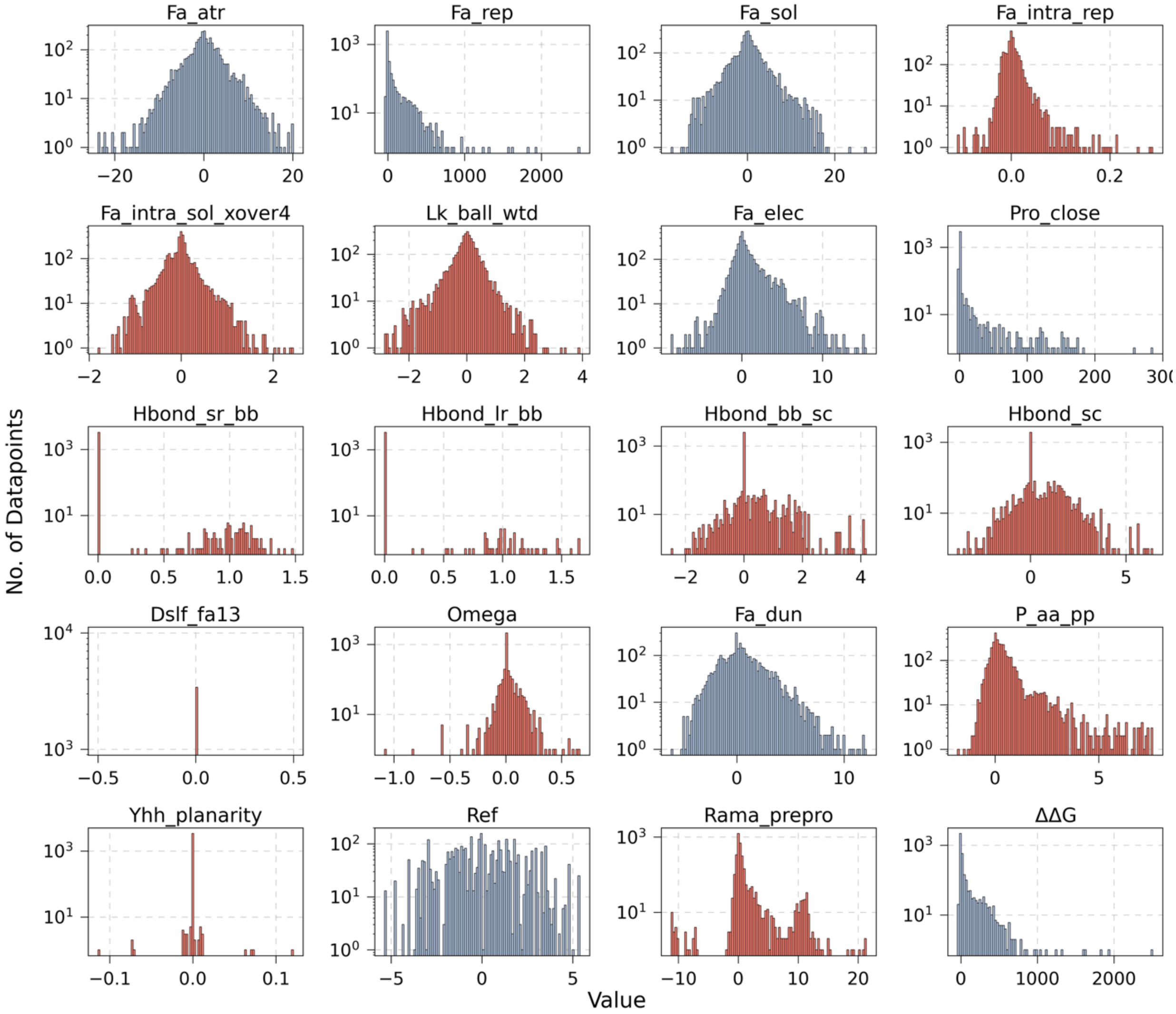


**Figure S13.** Distribution of all the 19 Rosetta terms and ΔΔG energies. The blue bar plots indicate term used as the input features and red bar plots indicate terms discarded as the input features.

**Table S1.** Possible input features. The features are divided into four categories: (1) One-hot encoding of 21 variables representing 20 amino acids and 1 stop codon; (2) AAIndex features consisting of 19 numerical values based on physicochemical properties [6]; (3) Rosetta energetics with 20 energy terms related to protein stability and folding, including attractive forces (fa_atr), van der Waals repulsive forces (fa_rep, fa_intra_rep), solvation energy (fa_sol, Fa_intra_sol_xover4, lk_ball_wtd), dielectric electrostatics (fa_elec), Proline ring closing energy (pro_close), disulfide statistical energies (dslf_fa13), and hydrogen bonding contributions (hbond_sr_bb, hbond_lr_bb, hbond_bb_sc, hbond_sc); and (4) the Root mean square fluctuation (RMSF) derived from atomistic simulations, capturing protein flexibility and represented by a single feature. Note that only 8 of the 20 Rosetta energy terms are selected to be used in training final VEPs reported in this work (highlighted in *italic*).

| Features | Description | Dimension |
|---|---|---|
| 1. One-hot encoding | Representing 20 amino acids and 1 stop codon | 21 |
| 2. AAIndex Features [6] | Numerical values corresponding to various physicochemical properties | 19 |
| 3. Rosetta Energetics [7]: | | 8 |
| *a. fa_atr* | Inter LJ attractive | |
| *b. fa_rep* | Inter LJ repulsive | |
| *c. fa_sol* | Lazaridis-Karplus solvation energy | |
| d. fa_intra_rep | Intra LJ repulsion | |
| e. fa_intra_sol_xover4 | Intra residue LK solvation | |
| f. lk_ball_wtd | Weighted sum of anisotropic and LK solvation | |
| *g. fa_elec* | Columbic electrostatic potential | |
| *h. pro_close* | Proline ring closure energy | |
| i. hbond_sr_bb | Backbone-backbone HBond of close residues | |
| j. hbond_lr_bb | Backbone-backbone HBond of distant residues | |
| k. hbond_bb_sc | Sidechain-backbone HBond energy | |
| l. hbond_sc | Sidechain-sidechain HBond energy | |
| m. dslf_fa13 | Disulfide geometric potential | |
| n. omega | Backbone omega dihedral | |
| *o. fa_dun* | Dunbrack's statistics derived for internal energy of sidechain rotamers. | |
| p. p_aa_pp | Probability of amino acid at phi/psi | |
| q. yhh_planarity | Sinusoidal penalty on $\chi_3$ to prevent H-bonds formed in the phenol plane | |
| *r. ref* | Reference energy of each amino acid. | |
| s. rama_prepro | Backbone torsion preference term | |
| *t. ddG* | Total free energy change | |
| 4. RMSF | Root means square fluctuation of $C_\alpha$ atoms derived from atomistic simulations | 1 |

**Table S2** Hyperparameters used for training various models on five different protein datasets. The table lists the batch size (32, 64, 128, 1024, 2048), learning rate (0.005, $1\times10^{-3}$, $1\times10^{-4}$), number of dense layers (1, 2, 3), and number of nodes for NN models (100, 200). For CNN models, the number of convolutional layers and filter sizes are reported, while for GCN models, the graph thresholds (in Å) are specified. If mutational or positional splits used different hyperparameters, they are indicated separately within the corresponding cell.

| Model | avGFP | Bgl3 | GB1 | Pab1 | Ube4b |
|---|---|---|---|---|---|
| **LR** | Batchsize:128<br>Learning rate:$10^{-4}$ | Batchsize:128<br>Learning rate:$10^{-4}$<br>Mutational:<br>Batchsize: 1024 | Batchsize:128<br>Learning rate:$10^{-4}$ | Batchsize:128<br>Learning rate:$10^{-3}$ | Batchsize:64<br>Learning rate:$10^{-4}$ |
| **NN** | Batchsize: 32<br>Learning rate:$10^{-4}$<br>Dense layer: 3<br>Nodes: 100<br>Positional:<br>Batchsize: 64<br>Learning rate:5e-3<br>Dense layer: 1<br>Nodes: 1000 | Batchsize: 32<br>Learning rate:$10^{-4}$<br>Dense layer: 2<br>Nodes: 100 | Batchsize: 64<br>Learning rate:$10^{-4}$<br>Dense layer: 1<br>Nodes: 1000<br>Mutational:<br>Batchsize: 32<br>Dense layer: 3<br>Nodes: 100 | Batchsize: 128<br>Learning rate:$10^{-3}$<br>Dense layer: 3<br>Nodes: 100 | Batchsize: 64<br>Learning rate:$10^{-4}$<br>Dense layer: 3<br>Nodes: 100 |
| **CNN** | Batchsize: 64<br>Learning rate:$10^{-3}$<br>Convolutionlayer:5<br>Filter: 128<br>Kernal: [3, 1] | Batchsize: 64<br>Learning rate:$10^{-4}$<br>Convolutionlayer:1<br>Filter: 32<br>Kernal: [17, 1] | Batchsize: 32<br>Learning rate:$10^{-4}$<br>Convolutionlayer:3<br>Filter: 128<br>Kernal: [17, 1] | Batchsize: 128<br>Learning rate:$10^{-4}$<br>Convolutionlayer:3<br>Filter: 128<br>Kernal: [17, 1] | Batchsize: 128<br>Learning rate:$10^{-4}$<br>Convolutionlayer:5<br>Filter: 128<br>Kernal: [3, 1] |
| | Mutational:<br>Kernal: [17, 1]<br>Batchsize: 128 | Mutational:<br>Batchsize: 2048<br>Convolutionlayer:3<br>Filter: 128<br>Learning rate:$10^{-3}$ | Mutational:<br>Batchsize: 1024<br>Convolutionlayer:1<br>Kernal: [7, 1] | Mutational:<br>Batchsize: 64<br>Convolutionlayer:1<br>Kernal: [3, 1]<br>Learning rate:$10^{-3}$ | |
| | Positional:<br>Kernal: [17, 1]<br>Batchsize: 64 | Positional:<br>Convolutionlayer:5<br>Filter: 128<br>Learning rate:$10^{-3}$ | Positional:<br>Batchsize: 64<br>Convolutionlayer:1<br>Filter: 32<br>Kernal: [13, 1] | Positional:<br>Batchsize: 1024<br>Convolutionlayer:1<br>Filter: 32<br>Kernal: [7, 1] | Positional:<br>Batchsize: 32<br>Learning rate:$10^{-3}$<br>Convolutionlayer:1<br>Filter: 32<br>Kernal: [7, 1] |
| **GCN** | Batchsize: 32<br>Learning rate:$10^{-4}$<br>Convolutionlayer:2<br>Filter: 128<br>Graph threshold: 7 | Batchsize: 128<br>Learning rate:$10^{-4}$<br>Convolutionlayer:1<br>Filter: 32<br>Graph threshold: 6 | Batchsize: 32<br>Learning rate:$10^{-4}$<br>Convolutionlayer:5<br>Filter: 128<br>Graph threshold: 7 | Batchsize: 128<br>Learning rate:$10^{-4}$<br>Convolutionlayer:1<br>Filter: 32<br>Graph threshold: 7 | Batchsize: 128<br>Learning rate:$10^{-4}$<br>Convolutionlayer:3<br>Filter: 128<br>Graph threshold: 7 |
| | | Mutational:<br>Batchsize: 1024 | Mutational:<br>Batchsize: 64 | Mutational:<br>Batchsize: 64<br>Convolutionlayer:3<br>Filter: 128<br>Learning rate:$10^{-3}$ | Mutational:<br>Batchsize: 64<br>Convolutionlayer:1<br>Filter: 256 |
| | Positional:<br>Batchsize: 1024<br>Learning rate:$10^{-3}$<br>Convolutionlayer:1<br>Filter: 128 | | Positional:<br>Batchsize: 64 | Positional:<br>Batchsize: 1024<br>Filter: 256 | |

### SI References


1. Olson CA, Wu NC, Sun R. A Comprehensive Biophysical Description of Pairwise Epistasis throughout an Entire Protein Domain. Current Biology. 2014;24: 2643–2651. doi:10.1016/j.cub.2014.09.072

2. Melamed D, Young DL, Gamble CE, Miller CR, Fields S. Deep mutational scanning of an RRM domain of the Saccharomyces cerevisiae poly(A)-binding protein. RNA. 2013;19: 1537–1551. doi:10.1261/rna.040709.113

3. Starita LM, Pruneda JN, Lo RS, Fowler DM, Kim HJ, Hiatt JB, et al. Activity-enhancing mutations in an E3 ubiquitin ligase identified by high-throughput mutagenesis. Proceedings of the National Academy of Sciences. 2013;110: E1263–E1272. doi:10.1073/pnas.1303309110

4. Sarkisyan KS, Bolotin DA, Meer MV, Usmanova DR, Mishin AS, Sharonov GV, et al. Local fitness landscape of the green fluorescent protein. Nature. 2016;533: 397–401. doi:10.1038/nature17995

5. Romero PA, Tran TM, Abate AR. Dissecting enzyme function with microfluidic-based deep mutational scanning. Proceedings of the National Academy of Sciences. 2015;112: 7159–7164. doi:10.1073/pnas.1422285112

6. Gelman S, Fahlberg SA, Heinzelman P, Romero PA, Gitter A. Neural networks to learn protein sequence–function relationships from deep mutational scanning data. Proceedings of the National Academy of Sciences. 2021;118: e2104878118. doi:10.1073/pnas.2104878118

7. Alford RF, Leaver-Fay A, Jeliazkov JR, O'Meara MJ, DiMaio FP, Park H, et al. The Rosetta All-Atom Energy Function for Macromolecular Modeling and Design. J Chem Theory Comput. 2017;13: 3031–3048. doi:10.1021/acs.jctc.7b00125
